# Effects of psychedelics on neurogenesis and brain plasticity: A systematic review

**DOI:** 10.1101/2023.07.19.549676

**Authors:** Rafael Vitor Lima da Cruz, Richardson N. Leão, Thiago C. Moulin

## Abstract

In the mammalian brain, new neurons continue to be generated throughout life in a process known as adult neurogenesis. The role of adult-generated neurons has been broadly studied across laboratories, and mounting evidence suggests a strong link to the HPA axis and concomitant malfunctions in patients diagnosed with mood disorders. Psychedelic compounds, such as phenethylamines, tryptamines, cannabinoids, and a variety of ever-growing chemical categories, have emerged as therapeutic options for neuropsychiatric disorders, while numerous reports link their effects to increased adult neurogenesis. In this systematic review, we examine studies assessing neurogenesis or neurogenesis-associated brain plasticity after psychedelic interventions and aim to provide a comprehensive picture of how this vast category of compounds regulates the generation of new neurons. We conducted a literature search on PubMed and Science Direct databases, considering all articles published until January 31, 2023, and selected articles containing both the words “neurogenesis” and “psychedelics”. We analyzed experimental studies using either *in vivo* or *in vitro* models, employing classical or atypical psychedelics at all ontogenetic windows, as well as human studies referring to neurogenesis-associated plasticity. Of a total of 205 articles, 68 met all the necessary conditions for further review. Our findings were divided into five main categories of psychedelics: CB1 agonists, NMDA antagonists, harmala alkaloids, tryptamines, and entactogens. We described the outcomes of neurogenesis assessments and investigated related results on the effects of psychedelics on brain plasticity and behavior within our sample. In summary, this review presents an extensive study into how different psychedelics may affect the birth of new neurons and other brain-related processes. Such knowledge may be valuable for future research on novel therapeutic strategies for neuropsychiatric disorders.

## Introduction

According to the Global Burden of Diseases study, used by the World Health Organization (WHO) for strategic planning, 264 million people, or about 4.5% of the world population, suffer from Major Depressive disorder (MDD). It is recognized as one of the most debilitating illnesses on a global scale, substantially significantly affecting daily activities, quality of life, cognitive abilities, and work productivity (James et al., 2018). MDD can be defined as a sustained anhedonic state that can be continuous or episodic, strongly influences self-esteem and significantly impacts social, family and professional life (Lépine & Briley, 2011). Mood and anxiety disorders are the most prevalent mental illnesses and the third most prevalent cause of disability, contributing to the global burden of disease (WHO, 2012). The majority of pharmacological interventions aiming to treat mood disorders such as MDD are benzodiazepines or Selective Serotonin Reuptake Inhibitors (SSRIs). However, these classes of drugs do not elicit positive outcomes for about 50 to 60% of patients, leading to a condition characterized as treatment-resistant depression (TRD) (Nestler et al., 2002). SSRIs, the most modern class of antidepressants, are taken daily, with an onset of the desired effects close to one month after the beginning of treatment. However, these medications can trigger adverse effects that appear early on and last for the duration of the therapy. These drugs also have a high risk of being misused, as individuals undergoing treatment tend to become physically dependent or addicted, even with the accompanying lethargy induced by them (Wong and Licinio, 2001).

The pathophysiology of depression is not yet fully understood; however, empirical data from classical antidepressants have led to the widely accepted monoamine hypothesis, which predicts that this disorder arises from a deficiency or imbalance of monoamine neurotransmitters. It is worth noting that several studies provide support for this theory. First, standard antidepressants primarily operate on the monoamine neurochemical route, aiming to re-establishing dopamine (DA), noradrenaline (NA), and serotonin (5-HT) levels to homeostatic concentrations (Heninger et al., 1996); Second, monoamine antagonists like reserpine, typically used for arterial hypertension, can induce depressive symptoms when taken over extended periods (Baumeister et al., 2003; Freis, 1954; De Freitas et al., 2016); Third, treatments for MDD and anxiety disorders usually require chronic, daily dosages for at least a month to produce meaningful effects (Kempermann, 2002). The latter observation has also led to a reinterpretation of the long-standing monoamine hypothesis of depression to what is now termed the neurogenic hypothesis of depression. This revised theory suggests that depression correlates with a decrease in the formation of new neurons in the adult brain, a process that seems to be revived by prolonged antidepressant treatment (Jacobs et al., 2000).

Adult neurogenesis is the process by which new neurons are continuously added to the brain throughout the life of an organism. Neurogenesis seems to be ubiquitous to all species with a central nervous system (Barker et al., 2011), and for many of them, the process is confined to specific regions (Barnea and Pravosudov, 2011; Drew et al., 2013). In rodents, it is restricted to two zones: the olfactory bulb, driven by the neural stem cells (NSCs) located in the subventricular zone (SVZ), and the dentate gyrus sub-region of the hippocampus, driven by the radial glial-like cells (RGL) (Laplagne et al., 2006). The foundations of the neurogenic theory of depression are supported by empirical data from clinical and preclinical studies aimed at understanding how the neurogenesis process is reverted to homeostatic levels when SSRI chronic treatment is applied (Miller and Hen, 2015). However promising, alternative pathways to the proposed hypothesis are under discussion (Data-Franco et al., 2017; N. X. Li, Hu, Chen, & Zhang, 2022; Raphael Mechoulam & Parker, 2013; Sanches, Quevedo, & Soares, 2021; Yuan et al., 2015) and new biochemical routes to treat depression are emerging, including the induction of neurogenesis independent of direct 5-HT modulation (Idell et al., 2017; Reiche et al., 2018). Among the chemical candidates for novel antidepressants, encouraging results have been found with the use of psychedelics (Aleksandrova & Phillips, 2021; DeVos & Miller, 2013; Muttoni, Ardissino, & John, 2019).

Psychedelics act mainly through brain plasticity, initially changing neuronal functionality at the molecular level by producing electrophysiological changes that stimulate neurotrophic signaling (Browne & Lucki, 2013; Castrén, Voikar, & Rantamaki, 2007; Magaraggia, Kuiperes, & Schreiber, 2021; Muscat, Hartelius, Crouch, & Morin, 2021). Ultimately, they induce neurite growth (Numakawa et al., 2010; Saengsawang & Rasenick, 2016; V. B. B. Thompson et al., 2012), synaptic remodeling (R.-J. Liu et al., 2013; Zhou & Song, 2001), neurogenesis (García-Cabrerizo & García-Fuster, 2016; Lima da Cruz, Moulin, Petiz, & Leão, 2018; F. Liu et al., 2017), and oxidative stress reduction (Frecska, Bokor, & Winkelman, 2016; Frecska, Szabo, Winkelman, Luna, & McKenna, 2013; Szabo, 2015). Thus, it is believed that psychedelics can create a window of opportunity for therapists to introduce cognitive-behavioral treatment strategies and produce long-lasting effects, which are independent of the classical pharmacological approaches to treat the hypothesized neurotransmitter imbalance (Keeler et al., 2021; Nichols, 2016; Worrell and Gould, 2021). Such a holistic and personalized approach can better integrate patients into the treatment process, reducing the current disconnection between popular beliefs on mental illnesses and scientific-guided psychiatric interventions (Healy, 2004; Lacasse & Leo, 2005).

Despite the encouraging perspectives on the applications of psychedelics, their safe employment requires a deeper understanding of their mechanisms, as the currently available compounds generally target multiple neurotransmitter systems and may lead to undesired effects (Belouin & Henningfield, 2018; Brunton, Chabner, & Knollmann, 2011; Geyer, Nichols, & Vollenweider, 2009). Moreover, the effects on brain physiology are shown to depend on ontogeny (J. Liu et al., 2006; Riga, Bortolozzi, Campa, Artigas, & Celada, 2016; Skaper & Di Marzo, 2012), gender (Lee, Wainwright, Hill, Galea, & Gorzalka, 2014; Realini et al., 2011; Rubino et al., 2008), dose (Fortunato et al., 2009; Maeda et al., 2007; Marinova, Walitza, & Grünblatt, 2017) and chemical interactions (Canales & Ferrer-Donato, 2014; Zuo et al., 2018). For this reason, we sought to cover the effects of such compounds on the plasticity process associated with neurogenesis (Christie & Cameron, 2006; Kempermann, 2012). To categorize these compounds, we adapted a classification done elsewhere (Calvey & Howells, 2018). Finally, we discuss findings encompassing any effect on the molecular, cellular, physiological and behavioural levels reported for *in vivo* or *in vitro* models related to neurogenesis.

## Methods

### Search and inclusion criteria

We followed the Preferred Reporting Items for Systematic Reviews and Meta-analysis (PRISMA) guidelines to standardize the workflow (Liberati, Tetzlaff, & Altman, 2009). All relevant articles were selected during a search using the PubMed and Science Direct databases in January 2023, where we searched for the combination of the words “psychedelics” and “neurogenesis”, including their MeSH terms, without limitations regarding publication date. Titles and abstracts were first scanned for articles presenting original results involving the assessment of neurogenesis with the intervention of psychedelic compounds. Psychedelics were defined and classified as described in (Calvey & Howells, 2018). Articles not written in English were also considered, but none passed our inclusion criteria, leaving only articles written in English to our final sample. Posterior to the systematic search, a reference screening on all reviews gathered was performed. Experimental articles with titles containing any psychedelic drug were selected for abstract analysis under the same inclusion criteria. Selected articles underwent an elimination process, excluding duplicates, absent abstracts, conference notes, editorials, book chapters, and case reports. Articles that did not intend to test or discuss psychedelics in any organic model and did not report any effect of psychedelics on neurogenesis were removed, during the abstract screening phase, articles that used neuroplasticity biomarkers as an outcome measure were included even if they were not found by the initial search in the databases. Then, we sorted reports by the drug chemical group for further scrutiny. Reviews were not included in the analysis but were considered in the discussion of results. The PRISMA flowchart for article selection is depicted in **Figure 1**.

**Figure 1.**
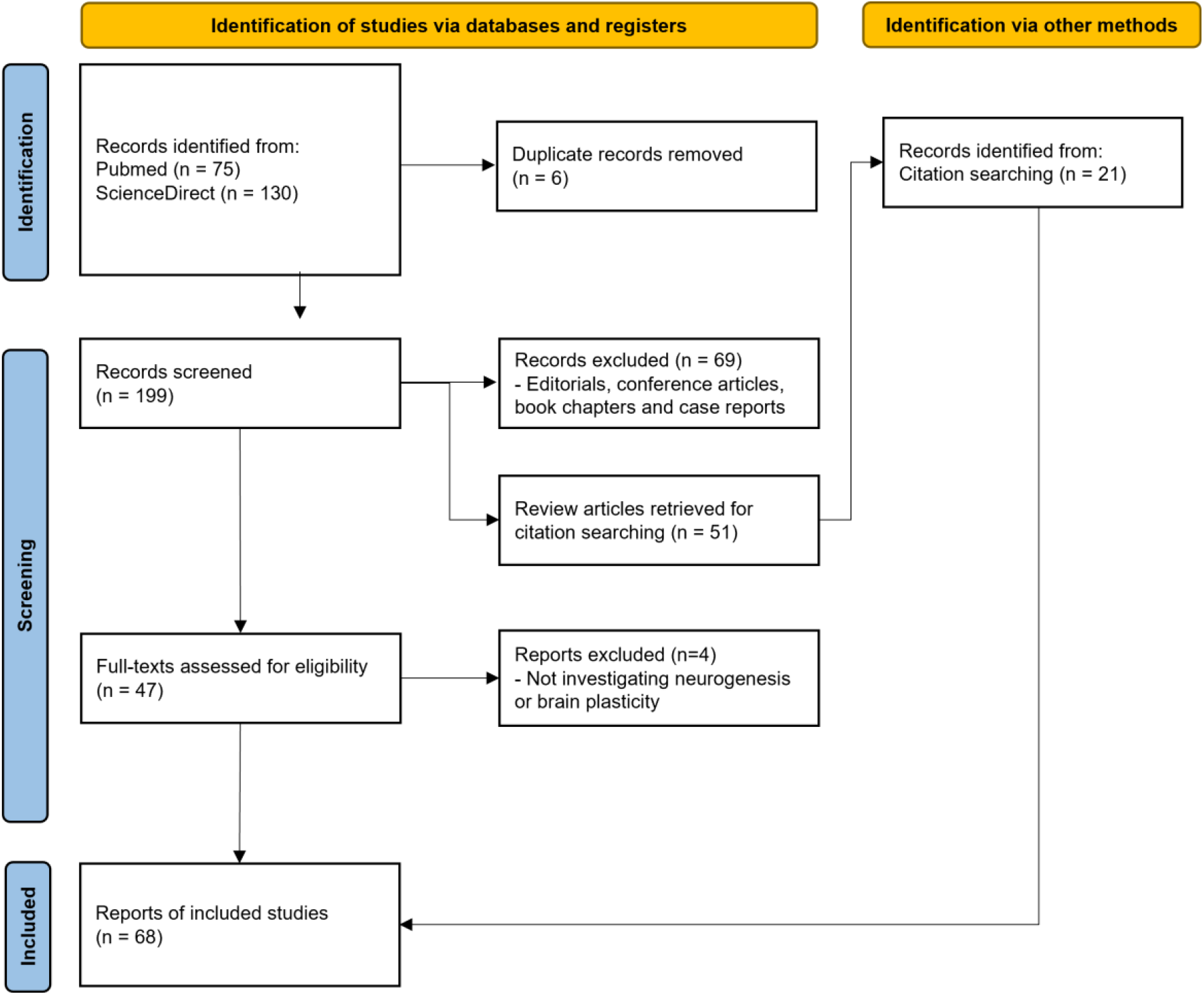
Flowchart of article selection following PRISMA guidelines. A total of 205 articles were initially identified from PubMed and Science Direct databases. After assessing abstracts, and conducting full-text analysis, 68 experimental articles were included in the final sample.

### Data extraction and analysis

Articles were organized in an Excel file by (1) article information, containing first and last author, author affiliation, journal, title, year of publication, and psychedelic chemical group; (2) methods information, such as drug dose and concentration, treatment regimen, experimental model of choice, techniques used, and (3) summary of results, highlighting outcome variables and reported effect on neurogenesis. For examination, a study location map (**Figure 2**) was generated using a web tool (www.mapinseconds.com), and an analysis of publications over the years (**Figure 3**) was performed using Microsoft Office Excel and GraphPad Prism.

**Figure 2.**
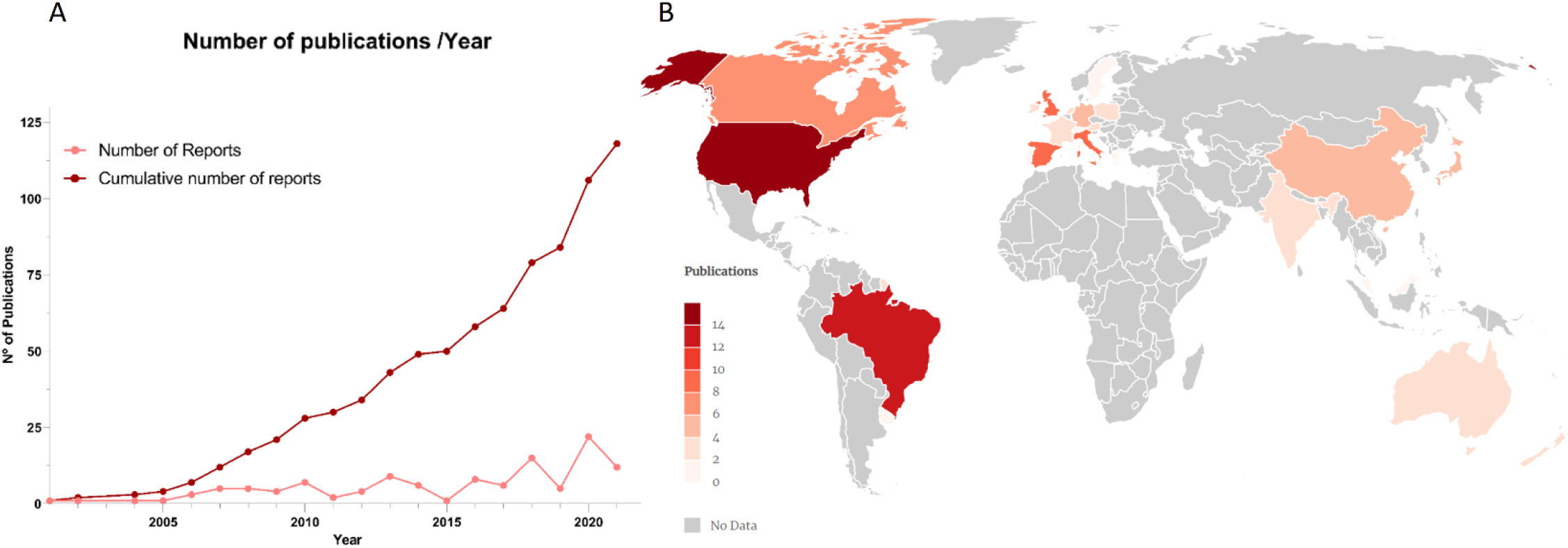
(A) Publication trend as cumulative (dark red) or absolute (light red) number for original articles and reviews identified by our search. (B) Localization of laboratories, identified by the affiliation of the correspondent author, which published original articles investigating the relationship between psychedelics and neurogenesis or neurogenesis-related plasticity (darker colours represent a higher number of publications, grey locations denote no publication record in our sample).

**Figure 2.**
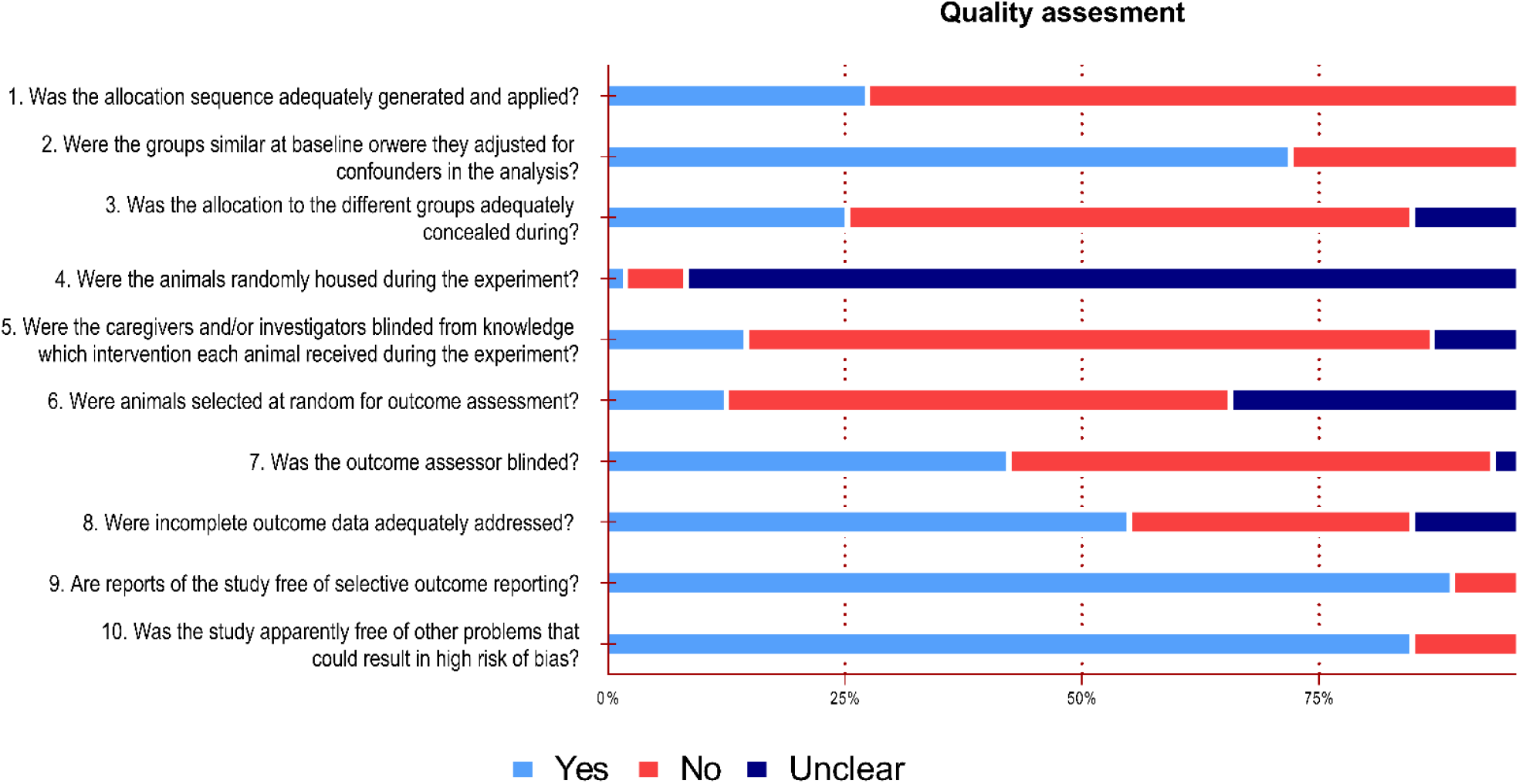
Results of SYRCLE Risk of Bias Quality assessment for the studies with animal experiments. Percentages of studies within a risk of bias assessment (Yes/ No/ Unclear) are represented in the x-axis for each risk of bias item.

**Figure 3.**
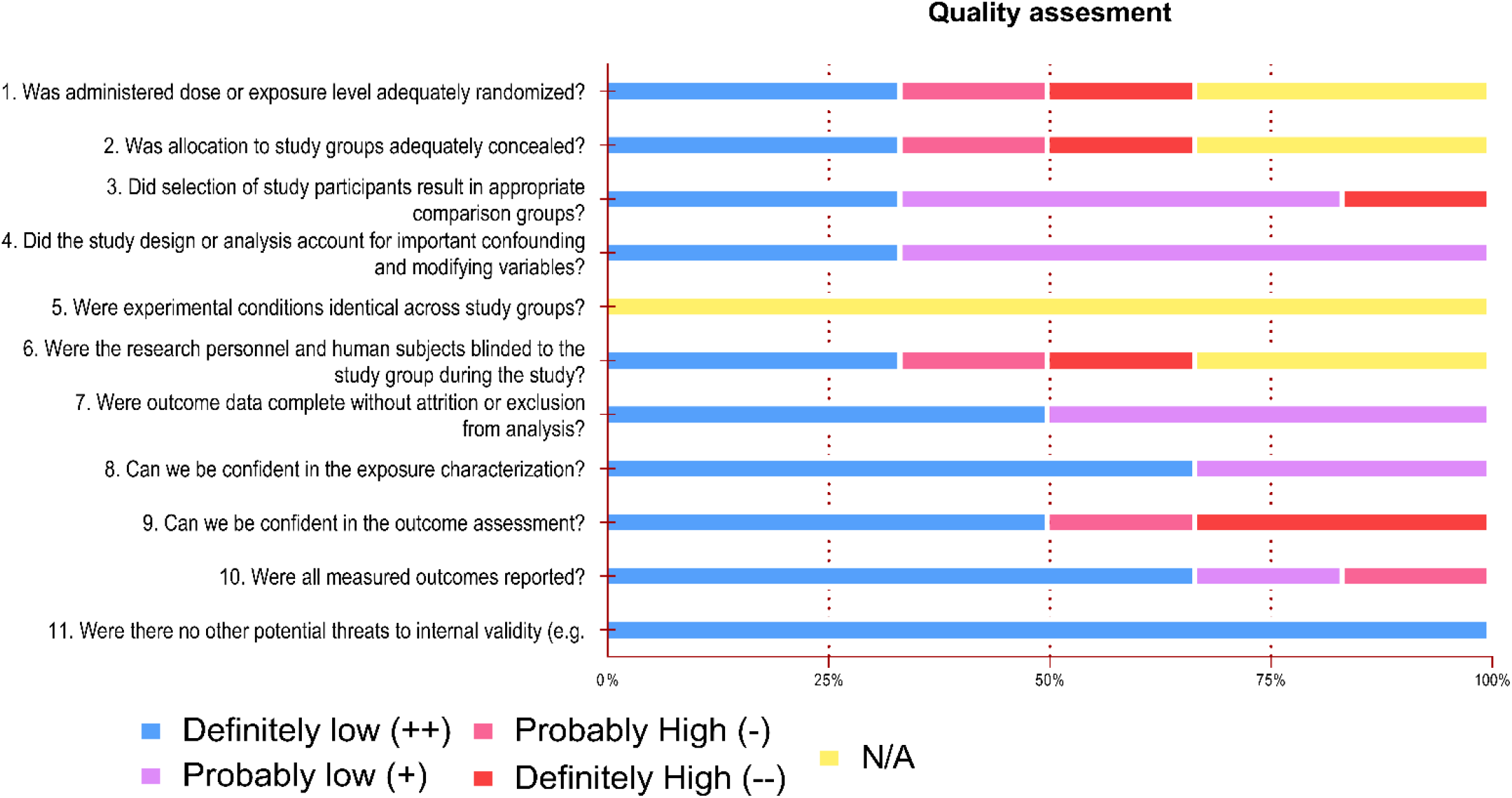
Results of OHAT Risk of Bias quality assessment of studies using human subjects. Percentages of studies within a risk of bias level are represented in the x-axis for each risk of bias item. Low risk of bias (blue, purple), unclear risk of bias (yellow), and high risk of bias (pink, red).

### Methodological quality assessment

We performed a qualitative analysis of the *in vivo*, animal and human studies, applying the criteria from SYRCLE (Hooijmans et al., 2014) and OHAT (OHAT, 2015) risk of bias (RoB) assessment tools, respectively. Both tools focus on methodological application, randomization of subjects, experimenter blinding, outcome measurements and reporting of methods. The results are reported in **Figure 2** for human studies and **Figure** for *in vivo* studies. Excel tables were generated for all categories, summarizing the method and results. *In vitro* studies were not analyzed for risk of bias. Considering the purpose of this review, electrophysiological assessment of acute slices was considered *ex vivo* and thus analyzed with the animal studies via SYRCLE RoB. However, long-term cultured neurons, even when removed from embryos, were not considered *ex vivo* but *in vitro*. Articles containing in vitro and in vivo experiments report their results independently. Finally, for the first criterion of SYRCLE RoB, “*Was the allocation sequence adequately generated and applied?*”, due to the poor reporting in this category, we graded positively articles that described any level of randomization. Moreover, for the second criterion of the SYRCLE RoB “*Were the groups similar at baseline, or were they adjusted for confounders in the analysis?”* for a study to be classified positively, we deemed necessary the reporting of age, gender and, when applicable, disease model onset.

## Results and Discussion

### Article search and inclusion

For this systematic review, we followed the PRISMA recommendations (Liberati et al., 2009), and included articles from two databases (PubMed and Science Direct) to identify articles related to the effects of psychedelics on neurogenesis and neurogenesis-related neuronal plasticity. A total of 205 articles were initially identified, 75 from PubMed and 130 from Science Direct. First, we assessed the abstracts of selected articles, removed conference notes, editorials, book chapters, and case reports (n=69), and excluded 32 publications that did not test or discuss the effects of psychedelics on neurogenesis or neuronal plasticity. Four additional articles were excluded after a full-text analysis for the same reasons. Lastly, narrative reviews were screened for further article selection, following the abovementioned criteria, resulting in 21 additional references. Our final sample contained 68 experimental articles examining the effects of psychedelics on neurogenesis or related brain plasticity outcomes. Of these, 6 were conducted in humans, 44 *in vivo*, and 11 *in vitro*. Additionally, 7 of the studies used both in *vivo* and in *vitro* approaches. Following article inclusion, we extracted metadata such as location, year of publication, and authorship, in addition to the methodological information and summarization of results. **Figure 1** summarizes the searching and selection process.

### Publication trends

The oldest articles in our sample date back to 2001, indicating that investigations on the effects of psychedelics on brain plasticity are a rather recent pursuit of the neuroscientific field. Neurogenesis in adult mammals was only described in 1977 (Kaplan & Hinds, 1977), and the process was not widely accepted as occurring in humans until 1998 (Eriksson et al., 1998) – although still controversial for part of the neuroscience community (Sorrells et al., 2018). Moreover, the prohibition of psychedelics by the UN Act of 1971 may have contributed to delaying research on these compounds (Ninnemann, Stuart, & Andersen, 2011; Rucker, Iliff, & Nutt, 2018). Nevertheless, it is worth highlighting that attention towards psychedelics has experienced a substantial increase during the past two decades (**Figure 2A**), with the most prolific publication years being 2013 (8 original articles; 2 reviews), 2018 (9 experimental articles; 6 reviews), and 2020 (8 experimental articles; 14 reviews).

Lastly, we can observe that most of the publications in our sample originated from a limited number of countries (**Figure 2B**), most of which have a well-documented history of high scientific productivity (King, 2004). The USA produced the highest number of studies (n=18), which primarily focused on the effects of ketamine (n=7) and cannabinoids (n=4).

Notably, Brazil deviates from this trend as the only emerging economy displaying a substantial volume of publications in the field of psychedelic research. This may be explained by the legal status of the consumption of the *ayahuasca* brew, allowed for religious practices (Labate & Feeney, 2012; Svobodny, 2014), as it is a traditional drink consumed ritualistically by indigenous people in the Amazonian basin (Labate & Cavnar, 2013). Accordingly, most Brazilian articles reviewed investigate the *ayahuasca* concoction or its main components, such as β-carbolines and tryptamines (n=8).

### Sample description and risk of bias assessment

A summary of methodological descriptions from the studies can be found in **Supp. Tables 1-6**. The main findings and discussion points of each of these studies are outlined in **Tables 1-6** and the following sections of this article.

**Table 1.**
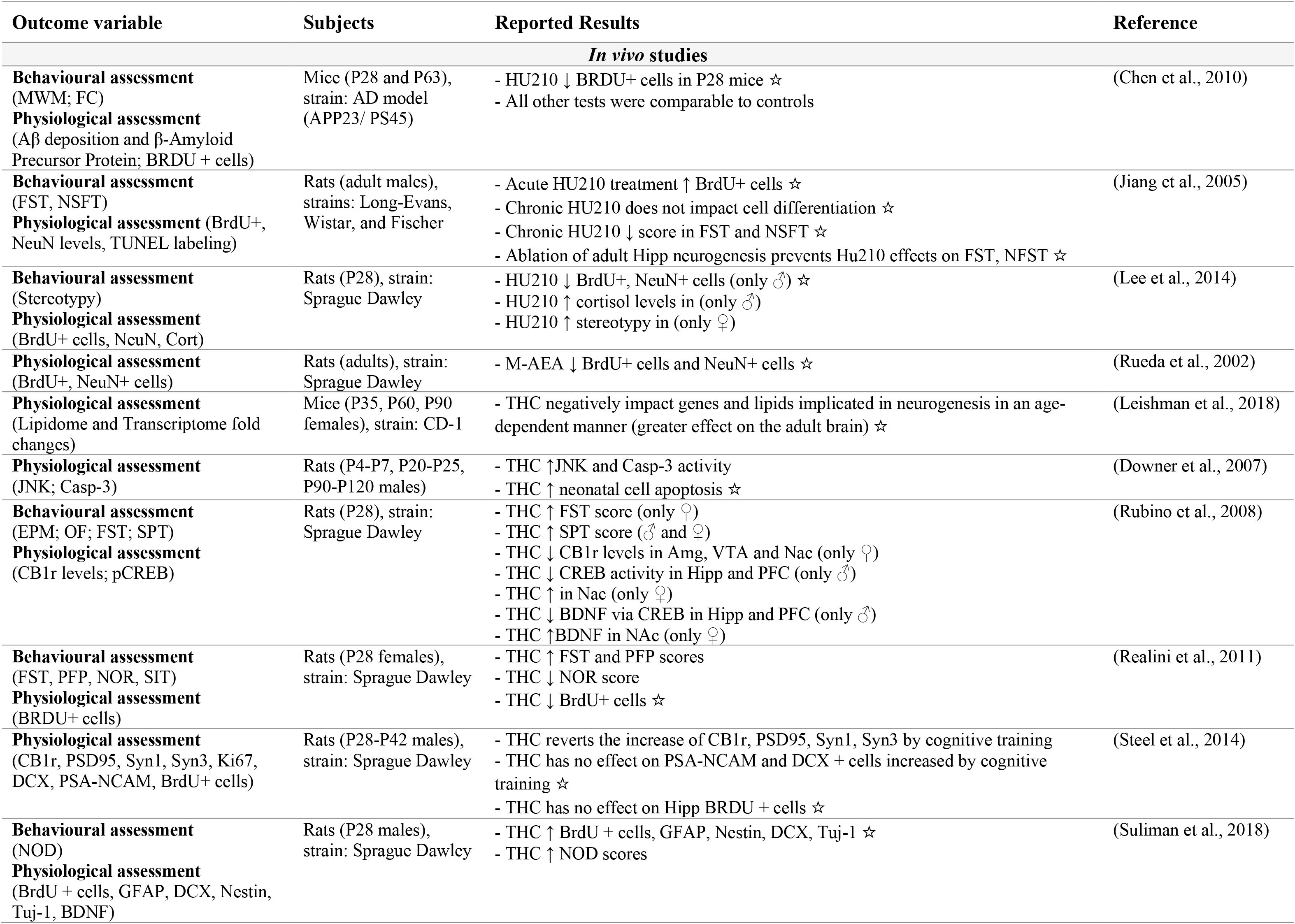
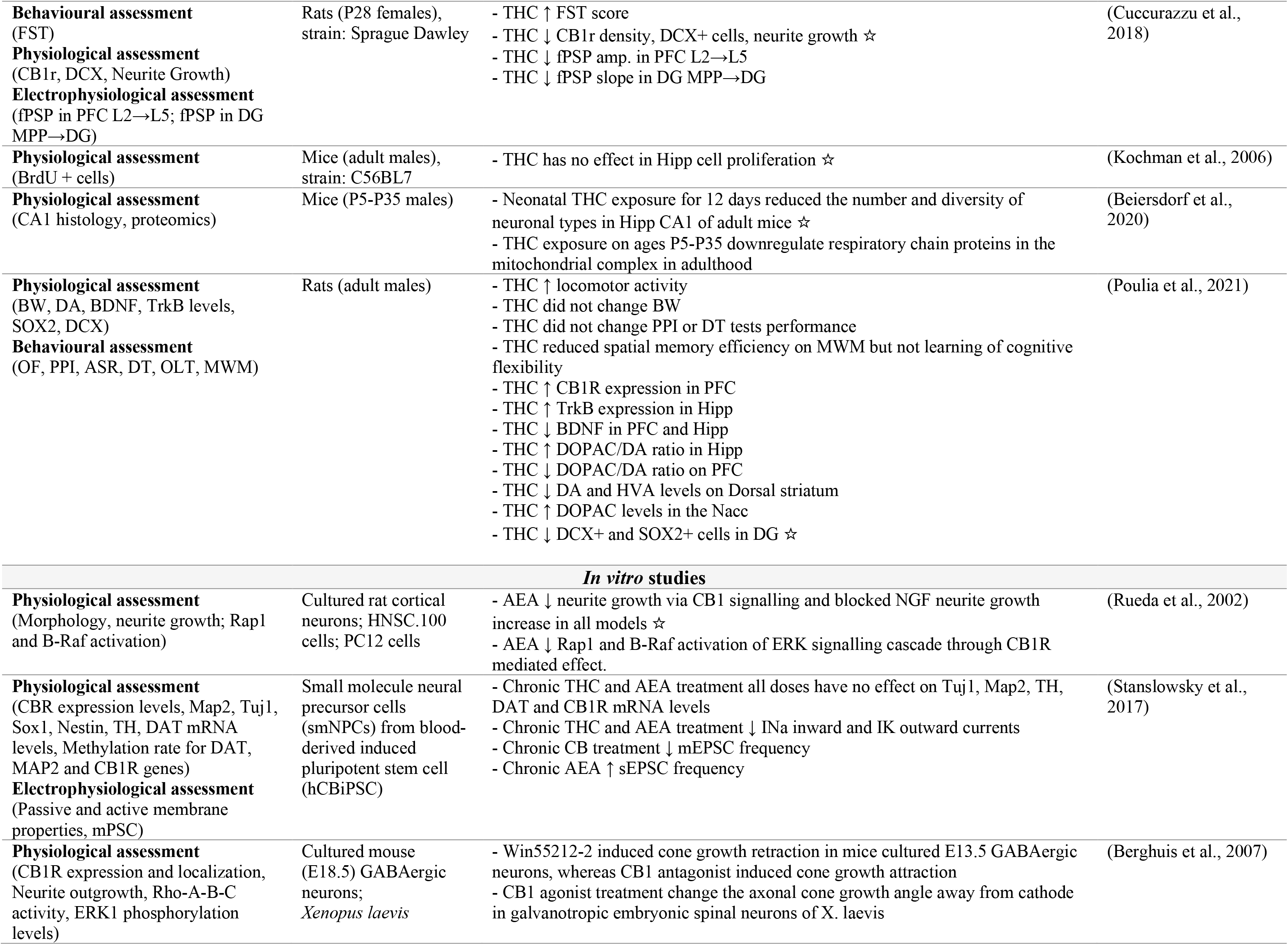
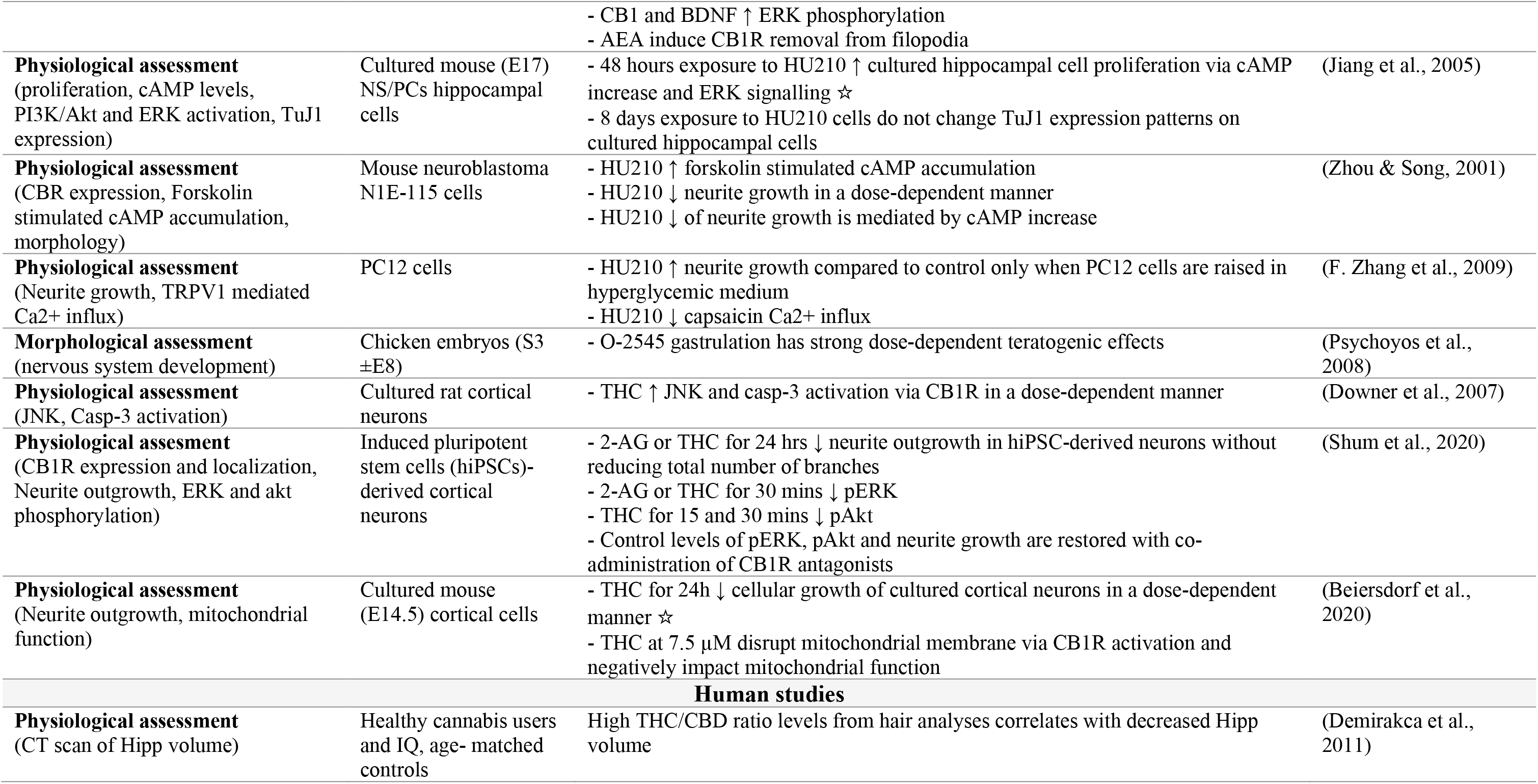
Results reported by CB1 agonists studies. ☆ specifies outcomes directly indicating neurogenesis through assessment of cell proliferation or differentiation. Abbreviations: MWM (Morris Water Maze); FC (Fear Conditioning); BrdU (Bromodeoxyuridine); FST (Forced Swim Test); NFST (Novelty suppressed Feeding Test); NOD (Novel Object Discrimination); cort (corticosterone); NeuN (Hexaribonucleotide Binding Protein-3); DCX (doublecortin); Tuj-1 (Neuron-specific class III beta-tubulin); M-AEA (Methanamide); JNK (c-Jun N-terminal kinases); Casp-3 (caspase 3); fPSP (Field Post-Synaptic Potential); mEPSC (miniature spontaneous Excitatory Post Synaptic Currents); DG (Dentate Gyrus); VTA (Ventral Tegmental Area); Hipp (Hippocampus); MPP (Medial Perforant Path); Rap1 (Ras-proximate-1 or Ras-related protein 1); B-raf (serine/threonine-protein kinase B-Raf); ERK (extracellular signal-regulated kinase); PPI (Pre-pulse Inhibition); ASR(Acoustic Startle reflex); DT (Discrimination tests); OLT(Object location test); smNPCs (small molecule Neuronal Precursor Cells); Map2 (Microtubule-associated protein 2); sox1 (SRY-box transcription factor 1); Nestin (neuroepithelial stem cell protein); TH (Tyrosine Hydroxylase); DAT (Dopamine Transporter); Rho -A-B-C (Rho family of GTPAses Ras); PSA-NCAM (polysialiated neural cell adhesion molecule); ERK1 (extracellular signal-regulated kinases); ROCK (serine-threonine kinase Rho kinase); pCREB (phosphorylated cAMP response element-binding protein); syn1 (synapsin-1); syn3 (synapsin-3); PSD95 (Post-synaptic Density protein 95); NS/PC (Neural Stem Progenitor Cells); cAMP (cyclic Adenosine monophosphate); PI3K/Akt (phosphatidylinositol 3-kinases/Protein Kinase B); N1E-115 (Mouse Neuroblastoma cell line); TRPV1 (transient receptor potential cation channel subfamily V member 1); PC12 (Rat adrenal medulla pheochromocytoma derived cell line);hiPSC (human induced Pluripotent Steam Cell; DAGLA (Diacylglycerol lipase); FAAH (Fatty acid amide hydrolase); MGLL (Monoacylglycerol lipase); CT (Computerized tomography).

**Table 2.** Results reported by the reviewed NMDA antagonists’ studies. ☆ specifies outcomes directly indicating neurogenesis through assessment of cell proliferation or differentiation. Abbreviations: Nac (Nucleus accumbens); PFC (Prefrontal Cortex); Glu (Glutamate); DA (Dopamine); PV (Parvalbumin); 8-OHdG (8-Oxo-2’-deoxyguanosine); NOX1 (NADPH oxidase 1); NOX2 (NADPH oxidase 2); syn (synapsin); OF (Open Field Test); FST (Forced Swim Test); cit-c (cytochrome complex); Casp3 (Caspase 3); Creb (cAMP response element-binding protein); mTor (mechanistic target of rapamycin); GSK-3β (Glycogen synthase kinase 3 beta); sEPSC (spontaneous Excitatory Post-synaptic Current); pyr (pyramidal neuron); abGC (adult born Granule Cells); GCL (Granule Cell Layer); RGL (Radial-Glia Like Cells); NBQX (2,3-dioxo-6-nitro-7-sulfamoyl-benzo[f]quinoxaline); ERK (extracellular signal-regulated kinase); BrdU (Bromodeoxyuridine); NOR (Novel Object Recognition Test); BDNF (Brain Derived Neurotrophic Factor); MWM (Morris Water Maze Test); BW (Body Weight); PCNA (Proliferating cell nuclear antigen); PAG (Phosphate activated Glutaminase); Pax6 (Paired box protein Pax-6); Tbr2 (T-box brain protein 2); Notch2 (Neurogenic locus notch homolog protein 2); Ntn1 (Netrin-1); PPI (Pre-Pulse inhibition Test); PSD95 (Post-synaptic Density Protein 95); Akt (Protein Kinase B); NRG1 (Neuregulin 1); PANSS (Positive and Negative Syndrome Scale); TNF-α (Tumor necrosis factor-α); IL-6 (Interleukin 6); IL -18 (interferon-gamma inducing factor); HMS (Hood Mysticism Scale); CADSS (Clinician Administered Dissociative States Scale); NDES (Near-Death Experience Scale); HDRS (Hamilton Depression Rating Scale); NGF (Nerve Growth Factor); NTF3 (neurotrophin-3); NTF4 (neurotrophin-4); GDNF (glial-derived neurotrophic factor); SWA (Slow-Wave activity); MADRS (Montgomery–Åsberg Depression Rating Scale).

| Outcome variable | Subjects | Reported Results | Reference |
| --- | --- | --- | --- |
| *In vivo studies* |  |  |  |
| **Physiological assessment**<br>(Glu, Glutamine, GABA, and PV levels; NAc DA levels and DA turnover; 8-OHdG, NOX1, NOX2) | Mice (P11 and P70 males), strain: C57BL6 | - Ketamine ↑ PFC Glu and Glutamine levels (P11)<br>- Ketamine ↑ PFC Glu and GABA levels (P70)<br>- Ketamine ↓ PFC PV, NOX1 levels (P11 & P70)<br>- Ketamine ↓ Nac DA levels (P11 & P70)<br>- Ketamine ↑ NAc DA turnover (P11 & P70)<br>- Ketamine ↑ Nac and PFC, 8-OHdG and NOX2 levels (P11 & P70)<br>- Ketamine ↓ NAc NOX1 (P11)<br>- Ketamine ↑ NAc NOX1 (P70) | (Schiavone et al., 2020) |
| **Behavioral assessment**<br>(OF)<br>**Physiological assessment**<br>(Cell morphology, Syn, Glu, DA, Cit-C, Casp3 and Creb pathway protein levels) | Rats (P35-42 males), strain: Sprague Dawley | - Ketamine ↑ locomotor activity, ataxia and stereotypy<br>- Ketamine ↑ cell morphology abnormalities in CA1, CA2 and CA3<br>- Ketamine ↓ Syn levels in Hipp<br>- Ketamine ↑ DA, Glu, Cit-C and Casp3 levels on the Cortex and Hipp<br>- Ketamine ↓ CREB pathway protein levels. | (Zuo et al., 2018) |
| **Behavioral assessment**<br>(FST)<br>**Physiological assessment**<br>(mTOR and GSK-3β activity, synaptogenesis)<br>**Electrophysiological assessment**<br>(sEPSC) | Rats (adult males), strain: Sprague Dawley | - Ketamine + lithium ↑ mTOR signaling<br>- Ketamine and Ketamine + lithium ↑ sEPSC frequency and synaptogenesis;<br>- Ketamine and Ketamine + lithium ↓ score in FST.<br>- Ketamine ↑ neurite outgrowth | (R.-J. Liu et al., 2013) |
| **Physiological assessment**<br>(BrdU+ cells) | Rats (P56 males), strain: Sprague Dawley | - Ketamine ↑ BrdU+ cells after 2 and 3 weeks ☆<br>- Ketamine ↑ cell survival | (Keilhoff et al., 2004) |
| **Behavioral assessment**<br>(NOR)<br>**Physiological assessment**<br>(BrdU+ cells, BDNF levels) | Rats (P70-95 males), strain: Sprague Dawley | - Ketamine after training ↓ score in NOR<br>- Ketamine ↓ BDNF levels increase after cognitive training in NOR. | (Goulart et al., 2010) |
| **Behavioral assessment**<br>(FST, NSFT)<br>**Physiological assessment**<br>(mTOR signaling, Neurite outgrowth)<br>**Electrophysiological assessment**<br>(sEPSC) | Rats (adult males), strain: Sprague Dawley | - Ketamine ↑ mTOR signaling in the PFC<br>- Ketamine ↑ synaptogenesis and sEPSC in PFC<br>- Ketamine-induced increases in mTOR signaling and synaptogenesis is abolished by pre-treatment with NBQX or ERK blockers<br>- Ketamine ↓ score in FST and NFST;<br>- Ketamine-induced reductions in FST and NFST scores are abolished by pre-treatment with rapamycin, and ERK blockers | (N. Li et al., 2010) |
| **Behavioral assessment**<br>(OF, FST, SPT)<br>**Physiological assessment**<br>(BDNF, Syn and d-mTOR levels) | Rats (adult males), strain: Wistar | - Ketamine ↓ FST score<br>- Ketamine ↑ SPT score<br>- Ketamine ↑ BDNF, Syn and d-mTOR levels | (Akinfiresoye & Tizabi, 2013) |
| **Behavioral assessment**<br>(FST)<br>**Physiological assessment**<br>(BDNF Levels) | Rats (adult males), strain: Sprague Dawley | - mPFC BDNF block prevent ketamine-induced FST score reduction<br>- L-Type channel block prevents ketamine-induced FST score reduction | (Lepack et al., 2015) |
| **Behavioral assessment**<br>(MWM)<br>**Physiological assessment**<br>(BrdU+ cells) | Rats (P7) | - Ketamine ↓ BrdU+ cells in proliferation essay ☆<br>- Ketamine ↑ BrdU+ neurons in differentiation essay ☆<br>- Ketamine ↓ BrdU+ astrocytes in differentiation essay<br>- Ketamine did not change the apoptosis rate in the DG<br>- Ketamine ↓ the migration of newborn granule cells<br>- Ketamine ↓ RGL growth<br>- P7 exposure to Ketamine ↓ score in MWM when adults | (Huang et al., 2016) |
| **Physiological assessment**<br>(Neurite outgrowth, Spine density) | Rats (P56 females),<br>strain: Sprague Dawley | - Ketamine ↑ spinogenesis in rats Pyr PFC neurons. | (Ly et al., 2018) |
| **Morphological assessment**<br>(BW)<br>**Physiological assessment**<br>(PFC cell density and number,<br>PFC cell migration, PFC,<br>PCNA+, PAG+, Pax6+, Tbr2+,<br>BrdU+ cells, expression levels of<br>Notch2 and Ntn1)<br>**Behavioral assessment**<br>(NOR, FST, PPI) | Mice (P1, P7 and P56<br>males), strain: CD-1 | - PCP ↓ BW (P1)<br>- PCP ↓ NOR and PPI score<br>- PCP ↑ FST score;<br>- PCP ↓ PFC cell density and PAG+ cells (P56)<br>- PCP ↓ PFC BrdU+ cells (P7) ☆<br>- PCP ↓ PCNA+, Pax6+, Tbr2+ cells<br>- PCP ↓ expression levels of Notch2 and ntn1. | (Toriumi et al., 2012) |
| **Physiological assessment**<br>(BrdU+ cells) | Rats (adult males),<br>strain: Sprague Dawley | - Chronic PCP ↓ BrdU + cells ☆<br>- Chronic PCP did not alter survival or differentiation of abGC | (J. Liu et al., 2006) |
| **Behavioral assessment**<br>(Locomotion)<br>**Physiological assessment**<br>(BrdU+ cells) | Rats (P21 males), strain:<br>Sprague Dawley | - PCP ↑ BrdU+ cells ☆<br>- PCP ↓ locomotor activity. | (Tanimura et al., 2009) |
| **Physiological assessment**<br>(BrdU+ cells) | Mice (P56 males), strain:<br>CD-1 | - PCP 3, 10 mg/kg ↓ BrdU+ cells ☆ | (Maeda et al., 2007) |
| ***In vitro studies*** |  |  |  |
| **Physiological assessment**<br>(Neurite growth; Synaptophysin,<br>PSD95 and BDNF levels; Akt and<br>GSK3β phosphorylation levels;<br>NRG1 activation levels) | Cultured mice PFC<br>neurons | - PCP ↓ neurite growth<br>- PCP ↓ Synaptophysin and PSD95 levels<br>- PCP ↓ phosphorylation of Akt and GSK3β without affecting the expression levels<br>- PCP reduction of neurite growth and Akt GSK3β phosphorylation is dependent of NRG1 activation<br>- PCP ↓ BDNF levels, and this reduction is NRG1 activation dependent<br>- PCP ↓ NRG1 expression levels | (Q. Zhang et al., 2016) |
| ***Human studies*** |  |  |  |
| **Survey assessment**<br>(PANSS)<br>**Physiological assessment**<br>(TNF-α, IL-6 and IL-18) | Hospitalized ketamine-addicted and matching<br>healthy control subjects | - Ketamine-addicted subjects have ↑ IL-6, IL-8 levels and PANSS scores;<br>- Ketamine-addicted subjects have ↓ TNF-α levels | (Fan et al., 2015) |
| **Survey assessment**<br>(HMS, self-reports on cocaine craving and self-administration) | Cocaine-addicted subjects | - HMS correlates with overall improvement<br>- Ketamine treatment reduced cocaine self-administration | (Dakwar et al., 2018) |
| **Survey assessment**<br>(HDRS)<br>**Physiological assessment**<br>(Serum BDNF, NGF, NTF3, NTF4 and GDNF) | Bipolar patients within a depression episode and resistant to classical antidepressants | - BDNF levels in non-responders was reduced 7 and 14 days after treatment<br>- A single dose of ketamine reduced score in HDRS in all patients 14 days after treatment<br>- 48% of patients had a HDRS score $\leq 7$ points 14 days after treatment | (Rybakowski et al., 2013) |
| **Survey assessment**<br>(MADRS)<br>**Physiological assessment**<br>(Serum BDNF)<br>**Electrophysiological assessment**<br>(SWA) | Treatment-resistant MDD-diagnosed subjects | - Single ketamine dose ↓ MADRS score<br>- Single ketamine dose ↑ Serum BDNF levels 230 minutes after<br>- Single ketamine dose ↑ total sleep time, SWS and REM sleep<br>- SWA and BDNF levels increase positively correlates in responders | (Duncan et al., 2013) |
| **Survey assessment**<br>(MADRS)<br>**Physiological assessment**<br>(Serum BDNF) | Treatment-resistant MDD-diagnosed subjects | - Single ketamine dose ↓ MADRS score in ~50% of patients;<br>- BDNF increase negatively correlates with MADRS score;<br>- Single dose of ketamine ↑ BDNF serum levels in responders. | (Haile et al., 2014) |

**Table 3.** Results reported by the reviewed studies on Harmala alkaloids. ☆ specifies outcomes directly indicating neurogenesis through assessment of cell proliferation or differentiation. Abbreviations: FST (Forced Swim Test); OF (Open Field Test); BDNF (Brain Derived Neurotrophic Factor); SPT (Sucrose Preference Test); TST (Tail Suspension Test); PFC (Pre Frontal Cortex); Hipp (Hippocampus); EdU (5-Ethynyl-2’-deoxyuridine); GFAP (Glial Fibrillary Acidic Protein); Nestin (Neuroepithelial Stem Cell protein); Sox-2 (sex determining region Y-box 2); Tuj-1 (Neuron-specific class III beta-tubulin); Map-2 (Microtubule-associated protein 2); CNPase (2’,3’-Cyclic-nucleotide 3’-phosphodiesterase); Ki67 (Marker Of Proliferation Ki-67); SVZ (Subventricular Zone); Dentate gyrus of the Hippocampus (SGZ); NPC (Neural Progenitor Cell); PCNA (Proliferating cell nuclear antigen); CMS (Chronic Mild Stress); DCX (Doublecortin); SGZ (Subgranular Zone); GLT-1 (glutamate transporter 1);.

| Outcome variable | Subjects | Reported Results | Reference |
| --- | --- | --- | --- |
| *In vivo studies* |  |  |  |
| **Behavioral assessment**<br>(FST; OF)<br>**Physiological assessment**<br>(Hipp BDNF) | Rats (P60), strain: Wistar | - Harmine ↓ score in FST<br>- Harmine ↑ Hipp BDNF | (Fortunato et al., 2009) |
| **Behavioral assessment**<br>(FST; OF)<br>**Physiological assessment**<br>(Hipp BDNF). | Rats (P60), strain: Wistar | - Harmine ↓ FST score<br>- Harmine ↑ Hipp BDNF | (Fortunato et al., 2010a) |
| **Behavioral assessment**<br>(SPT, OFT)<br>**Physiological assessment**<br>(Adrenal Weight, blood ACTH and Hipp BDNF) | Rats (P60), strain: Wistar | - Harmine ↑ SPT score<br>- Harmine ↓ adrenal weight<br>- Harmine ↓ blood ACTH<br>- Harmine ↓ Hipp BDNF increase induced by CMS model | (Fortunato et al., 2010b) |
| **Behavioral assessment**<br>(TST; FST; SPT)<br>**Physiological assessment**<br>(Hipp and PFC BDNF; GLT-1; GFAP) | Mice (P56-P70 males), strain: C57BL6 | - Harmine ↓ TST, FST scores<br>- Harmine ↑ SPT score<br>- Harmine ↑ BDNF, DCX+ cells, GLT-1, GFAP ☆ | (F. Liu et al., 2017) |
| *In vitro studies* |  |  |  |
| **Physiological assessment**<br>(EdU+, Nestin, GFAP cells) | Human neural progenitor cells (hNPCs) | - Harmine ↑ proliferation and neuronal differentiation. | (Dakic, Maciel, et al., 2016) |
| **Physiological assessment**<br>(Musashi1, Nestin, Sox-2, Tuj-1, MAP-2, CNPase, GFAP levels, Ki67 +, DCX+ Cells) | Neurospheres from adult mice SGZ and SVZ | - Chronic B-Carbolines ↓ Musashi-1, Sox-2, Nestin levels in both NPC populations<br>- Chronic B-Carbolines ↑ neuronal number and size of adult neurospheres in both NPC populations ☆<br>- Chronic B-Carbolines ↑ ki67+ cells and PCNA levels<br>- Acute B-Carbolines ↑ Tuj1 and MAP-2 levels in both NPC populations, with exception to Harmine at SGZ for Tuj1<br>- Chronic Harmol and Harmaline ↑ CNPase levels at SVZ population<br>- Chronic B-Carboline ↑ GFAP in both NPC populations, with exception to Harmine at SGZ | (Morales-García et al., 2017b) |

**Table 4.** Results reported by the reviewed studies on psychoactive tryptamines. ☆ specifies outcomes directly indicating neurogenesis through assessment of cell proliferation or differentiation. Abbreviations: vDG (ventral Dentate Gyrus); sEPSC (spontaneous Excitatory Post Synaptic Current); Pyr (Pyramidal neurons); Nac (Nucleus accumbens); PFC (Prefrontal Cortex); SN (Substantia Nigra); Hipp (Hippocampus); VTA (Ventral Tegmental Area); CUS (Chronic Unpredictable Stress model); OF (Open Field); TST (Tail Suspension Test); FST (Forced Swim Test); EPM (Elevated Plus Maze); 5-HT (Serotonin); TPH-2 (Tryptophan Hydroxylase 2); ETC-1 (Mitochondrial electron transporter complex-I); DCX (doublecortin); GC (Granule Cells); BrdU (Bromodeoxyuridine); AHP (After Hyperpolarization); AP (Action Potential); LFP (Local Field Potential); TFC (Trace Fear Conditioning); AA (Amino acids) HTR (Head-Twitch Response); GDNF (Glial Cell Derived Neurotrophic Factor); BDNF (Brain Derived Neurotrophic Factor); NGF (Nerve Growth Factor); pet1 [pheochromocytoma 12 ETS (E26 transformation-specific)]; DRN (Dorsal Raphe Nucleus); MTT (3-[4,5-Dimethylthiazol-2-yl]-2,5-diphenyl tetrazolium bromide); Ki67 (Marker Of Proliferation Ki-67); AYA1 (santo Daime’s Ayahuasca decoction); CEB (*Banisteriopsis caapi* crude extract); CEP (*Psychotria viridis* crude extract); AYA2 (Ayahuasca prepared by authors); TrkB (Tropomyosin receptor kinase B); mTOR (mechanistic target of rapamycin).

| Outcome variable | Subjects | Reported Results | Reference |
| --- | --- | --- | --- |
| *In vivo studies* |  |  |  |
| **Physiological assessment**<br>(Neurite outgrowth, Spine density)<br>**Electrophysiological assessment**<br>(sEPSC) | Rats (P56 females), strain: Sprague-Dawley;<br>*Drosophila melanogaster* adults | - DOI, LSD ↑ Neurite outgrowth in *Drosophila* and Rats<br>- DMT and ketamine ↑ Spine density PFC Pyr neurons<br>- DMT ↑ sEPSC frequency and amplitude in PFC Pyr neurons | (Ly et al., 2018) |
| **Physiological assessment**<br>(BrdU+ cells, DCX+ cells, Neurite outgrowth)<br>**Electrophysiological assessment**<br>(Passive and active membrane properties) | Mice (P55-P70 males), strain: C57BL6 | - 5-MeO-DMT ↑ BrdU+ cells in vDG ☆<br>- 5-MeO-DMT ↑ DCX+ neurons in vDG ☆<br>- 5-MeO-DMT ↓ AHP and AP threshold in young GC on vDG<br>- 5-MeO-DMT ↑ maturation rate of young GC on vDG<br>- 5-MeO-DMT ↑ number of branches of young GC at vDG. | (Lima da Cruz et al., 2018) |
| **Behavioral assessment**<br>(OF)<br>**Electrophysiological assessment**<br>(LFP) | Mice (P120-P150 males), strain: C57BL6 | - 5-MeO-DMT ↓ anxiety-like behavior and electrophysiological LFP signature of anxiety | (Winne et al., 2020) |
| **Behavioral assessment**<br>(OF)<br>**Physiological assessment**<br>(Nac, PFC, VTA and SN mRNA and protein levels for GDNF, BDNF and NGF) | Rats (P60), strain: Wistar | - Ibogaine ↑ GDNF mRNA in VTA and SN 24h after injection;<br>- Ibogaine ↑ BDNF mRNA in all areas 24h after injection;<br>- Ibogaine ↓ BDNF mRNA in PFC 3h after injection<br>- Ibogaine ↑ NGF mRNA in all areas 24h after injection<br>- Ibogaine ↑ NGF mRNA in PFC and VTA 24h after injection<br>- Ibogaine ↑ GDNF protein levels in VTA 24h after injection<br>- Ibogaine ↑ proBDNF protein levels in Nac 24h after injection. | (Marton et al., 2019) |
| **Behavioral assessment**<br>(OF, FST, EPM)<br>**Physiological assessment**<br>(5-HT levels, TPH-2, Ca<sup>2</sup>ATPase and ETC-1 activity, 5-HT<sub>2C</sub>-R expression, Cell morphology) | Rats (P60), strain: Wistar | - MP ↑ 5-HT levels in Hipp and PFC in the CUS model<br>- MP ↑ TPH-2 activity in Hipp in the CUS model<br>- MP ↑ 5-HT<sub>2C</sub>-R expression in PFC on CUS and healthy animals<br>- MP ↑ 5-HT<sub>2C</sub>-R expression in Hipp on CUS but not on healthy animals<br>- MP ↑ Ca<sup>2</sup>ATPase activity in Hipp and PFC in the CUS model<br>- MP rescues ETC-1 downregulation induced by CUS<br>- MP ↓ anxiety levels and anhedonical phenotype in CUS animals<br>- MP ↓ cell morphological anomalies induced by CUS in Hipp and PFC | (Wankhar et al., 2020) |
| **Behavioral assessment**<br>(OF)<br>**Physiological assessment**<br>(BrdU+ cells)<br>**Electrophysiological assessment**<br>(Active membrane properties) | Mice (both sexes adults), strain: 5-HT<sub>2B</sub>-KO and *wt* control | BW723C86 ↑ firing rate on pet1+ DRN ex vivo<br>BW723C86 prevent 5-HT<sub>1A</sub> agonist induced inhibition of pet1+DRN in vivo<br>MDMA fail to ↑ locomotor activity on 5-HT<sub>2B</sub>-KO<br>Fluoxetine fails to ↑ BrdU+ cells in 5-HT<sub>2B</sub>-KO | (Belmer et al., 2018) |
| **Behavioral assessment** (TFC)<br>**Physiological assessment**<br>(BrdU+ cells) | Mice (adult males), strain: C57BL6 | Psilocybin do not change TFC score for acquisition;<br>Psilocybin ↑ fear response on trial 1 of TFC for extinction<br>Psilocybin ↓ fear response on trial 10 of TFC for extinction<br>Psilocybin ↓ BrdU+ neurons in DG ☆<br>NBOMe ↓ BrdU+ neurons in DG ☆ | (Catlow et al., 2013) |
| **Physiological assessment** (AA and monoamines levels, utilization rate of monoamines) | Rats (P60 males), strain: Wistar | Ayahuasca ↑ GABA levels in Hipp<br>Ayahuasca ↓ Glycine and GABA levels on Amygdala (dose dependent);<br>Ayahuasca ↑ 5-HT on Hipp;<br>Ayahuasca ↑ all monoamines on Amygdala<br>Ayahuasca 800mg/kg ↓ 5-HT use rate on Hipp<br>Ayahuasca ↓ monoamine consumption on amygdala | (Castro-Neto et al., 2013) |
| **Behavioral assessment** (Task oriented behavior; HTR) | Mice (P120-P150), strain: C57BL6;<br>Zebrafish larvae | DMT synthetic isoforms have comparable effect to its natural counterparts in a cue-oriented locomotion protocol<br>5-MeO-IsoDMT do not produce any HTR still have in vitro psychoplastogens properties | (Dunlap et al., 2020) |
| **Physiological assesment** (BrdU+ and DCX+ cells)<br>**Behavioral assesment** (MWM; NORT) | Mice (adult males), strain: C57BL6 | DMT ↑ survival and proliferation of newborn granule cells BrdU+ and DCX+ which is blocked by sigma-1R antagonist ☆<br>DMT ↓ escape latency in MWM is blocked by 5-HT2AR antagonist;<br>DMT ↑ exploration time in NORT. | (Morales-Garcia et al., 2020) |
| ***In vitro studies*** |  |  |  |
| **Physiological assessment** (cell viability by MTT and Calcein assay; ki67+ and PI+ cells) | SH-SY5Y cells | AYA1, CEB, CEP all doses ↑ cell viability with 48h incubation;<br>AYA2 1, 1.5 µg/mL ↑ cell viability with 72h incubation. | (Katchborian-Neto et al., 2020) |
| **Physiological assessments** (Neurite growth) | Cultured rat cortical neurons | DMT, isoDMT and 1-Me-DMT ↑ neurite growth in a dose comparable manner;<br>5-HT2A binding is necessary to neurite growth effect of all psychoplastogens tested. | (Dunlap et al., 2020) |
| **Physiological assessments** (Neurite growth, Spinogenesis) | Cultured rat cortical neurons | DOI, DMT, LSD ↑ Neurite growth and spinogenesis;<br>The effects of psychoplastogens are similar to increased BDNF in culture medium, and both act through TrkB and mTOR cascade as well 5-HT2A signaling | (Ly et al., 2018) |
| **Physiological assessment** (Musashi1, Nestin, Sox-2, PCNA, Tuj, Map-2, GADPH, GFAP, CNPase levels) | Cultured mice SGZ neurons | Chronic DMT ↓ of Musashi1, Nestin and SOX2 expression levels are blocked by Sigma-1R antagonist<br>Chronic DMT ↑ number of neurospheres and PCNA expression levels are blocked by sigma-1R antagonist;<br>Chronic DMT ↑ tuj, Map-2 GFAP and CNPase are blocked by sigma-1R antagonist | (Morales-Garcia et al., 2020) |

**Table 5.**
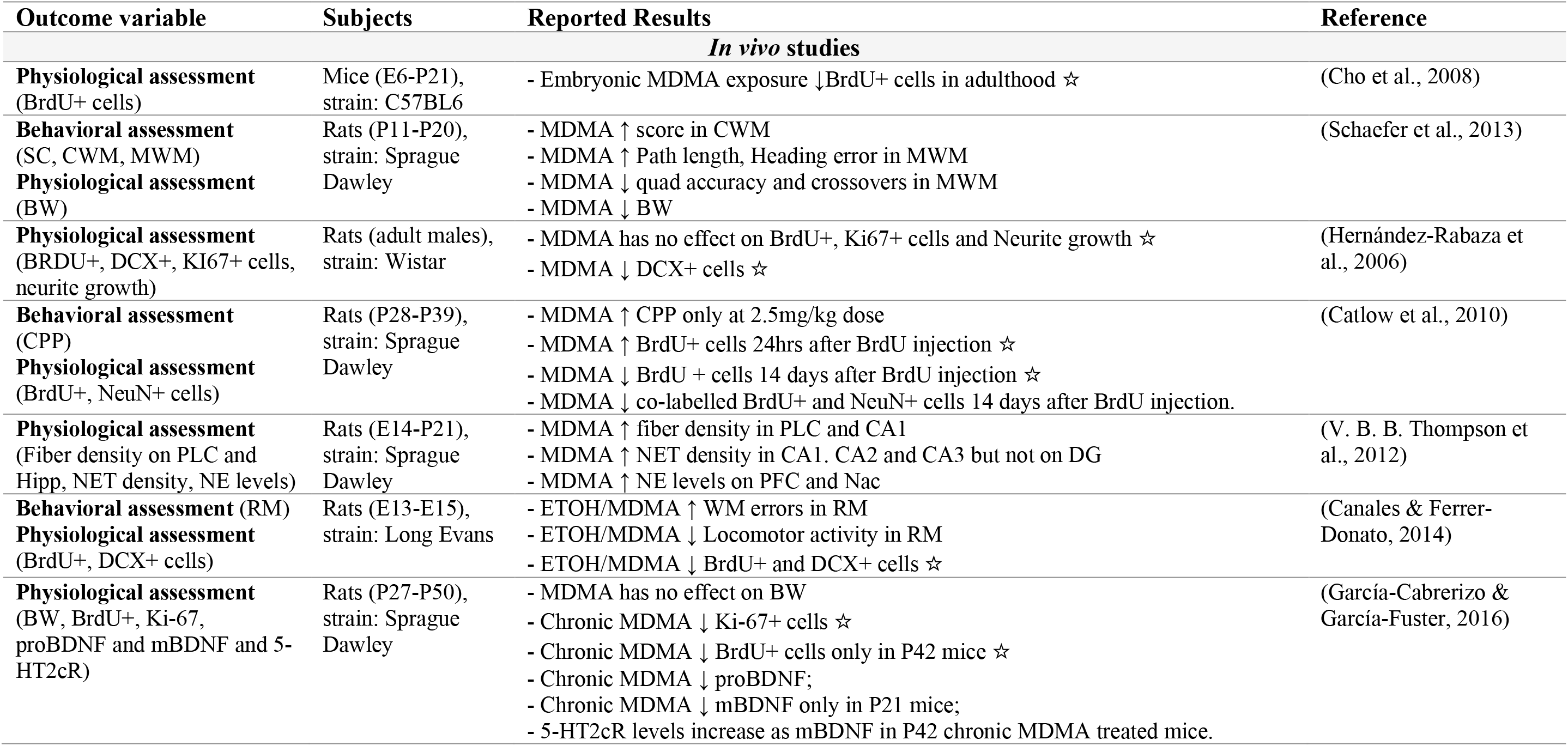
Results reported by the reviewed studies on Entactogens. ☆ specifies outcomes directly indicating neurogenesis through assessment of cell proliferation or differentiation. Abbreviations: SC (Straight Channel test); CWM (Cincinnati Water Maze); MWM (Morris Water Maze); BW (Body Weight); BrdU (Bromodeoxyuridine); DCX (Doublecortin); KI67 (Marker of Proliferation Ki-67); CPP (Conditioned Place Preference); PLC (Pre-limbic Cortex); Hipp (Hippocampus); NET (Noradrenergic Transporter); NE (Noradrenaline); DG (Dentate Gyrus); PFC (Prefrontal Cortex); Nac (Nucleus Accumbens); ETOH (Ethanol); RM (Radial Maze); proBDNF (precursor protein of BDNF); mBDNF (mature BDNF).

**Table 6.** Results reported by the reviewed studies on other psychedelics not covered by previous classifications. ☆ specifies outcomes directly indicating neurogenesis through assessment of cell proliferation or differentiation. Abbreviations: EPSC (Excitatory Post Synaptic Current); Pyr (Pyramidal); mPFC (medial Prefrontal Cortex); GluA2 (AMPA receptor subunit GluA2); LTD (Long term Depression); PKC (Protein Kinase C); BrdU (Bromodeoxyuridine); DG (Dentate Gyrus); PSD95 (Post-synaptic Density Protein 95); bassoon (Pre synaptic cytomatrix protein bassoon); MUPP1 (Multiple PDZ protein-1); Cos-7 (Fibroblast-like cell line from monkey kidney); PAK (P21-activated kinase); TrkA (Tropomyosin receptor kinase A); AA (Amino Acid); Tyr (Tyrosine).

| Outcome variable | Subjects | Reported Results | Reference |
| --- | --- | --- | --- |
| *In vivo studies* |  |  |  |
| **Electrophysiological assessment**<br>(Layer1 -↑ Layer 5 EPSC) | Mice (P14-P21), strain: 5-HT<sub>2A</sub>-KO and *w.t.* control | - DOI ↓ Layer 1 AMPA ESPC Amplitude in Layer 5 Pyr mPFC<br>- mPFC layer 5 Pyr of 5-HT<sub>2A</sub>-KO mice do not react to DOI treatment<br>- GluA2 subunit internalizations necessary for DOI-induced LTD in layer 5 Pyr mPFC<br>- PKC/ 5-HT<sub>2A</sub>R inhibitors block DOI-induced LTD in layer 5 Pyr mPFC | (Berthoux et al., 2019) |
| **Physiological assessment**<br>(BrdU + cells) | Rats (adult males), strain: Wistar | - DOI does not change BrdU+ cells on DG ☆<br>- Acute Ketanserin ↓ BrdU+ cells on DG ☆<br>- Chronic Ketanserin ↑ BrdU+ cells on DG ☆ | (Jha et al., 2008) |
| **Behavioral assessment**<br>(OF)<br>**Physiological assessment**<br>(BRdU+ cells)<br>**Electrophysiological assessment**<br>(Active membrane properties) | Mice (both sexes adults), strain: 5-HT<sub>2B</sub>-KO and *wt* control | - BW723C86 ↑ firing rate on pet1+ DRN *ex vivo*<br>- BW723C86 prevent 5-HT<sub>1A</sub> agonist-induced inhibition of pet1+DRN *in vivo*;<br>- DOI fail to ↑ HTR in 5-HT2B-KO;<br>- Fluoxetine fails to ↑ BRdU+ cells in 5-HT2B-KO. | (Belmer et al., 2018) |
| *In vitro studies* |  |  |  |
| **Physiological assessment**<br>(5-HT2AR, PSD95, Kalirin-7 and bassoon localization; MUPP1 activity; Dendritic spine remodeling; PAK phosphorylation levels) | Cultured rat cortical neurons | - DOI promotes spine shape remodelling without changing global spine density through PAK phosphorylation and subsequent Kalirin-7 activity | (Jones et al., 2009) |
| **Physiological assessment**<br>(TrkA tyr Phosphorylation levels at AA residue Tyr490 and Tyr785, Neurite Extension) | Human neuroblastoma SK-N-SH cells | - DOI all doses for 30 or 120 minutes ↑ TrkA phosphorylation in SK-N-SH cells<br>- DOI all doses for 30 or 120 minutes ↑ TrkA phosphorylation at AA residue Tyr490 but not Tyr785<br>- 5-HT2A and 5-HT2B block do not affect TrkA phosphorylation induced by DOI<br>- Chronic DOI 1,5 μM ↑ neurite extension (can be blocked by TrkA inhibitor) | (Marinova et al., 2017) |

#### Human studies

Of the 68 experimental articles identified in this study, 6 involved human participants. One case-control study examined the balance of CBD and THC phytocannabinoids in hair samples and their relationship to hippocampal mass (Demirakca et al., 2011). The other five studies explored the effects of ketamine in humans, including one cross-sectional study conducted with ketamine-addicted patients in Guangzhou, China (Fan et al., 2015), and four controlled trials (Dakwar et al., 2018; Duncan et al., 2013; Haile et al., 2014; Rybakowski, Permoda-Osip, Skibinska, Adamski, & Bartkowska-Sniatkowska, 2013). Quality assessment using the OHAT risk of bias tool is presented in **Figure 3**. Overall, the majority of articles showed ‘definitely low’ or ‘probably low’ risk of bias for most items. However, several articles had higher levels of bias or incomplete information for group randomization, allocation of subjects, and blinding. The criteria 5 is not applicable (NA) to the human studies included in our review. Moreover, half of the studies had subpar levels of outcome assessment confidence.

These results point to a diverse range of methodological robustness within the reviewed literature. The prevalence of low risk of bias in most articles suggests adherence to good research practices, implying a reliable foundation for drawing meaningful conclusions. Nonetheless, the identification of higher levels of bias in some studies and incomplete information in group randomization, allocation of subjects, and blinding, highlights specific areas for improvement in future research.

#### In vitro studies

Of the 18 original articles performing *in vitro* experiments, the majority utilized cannabinoid agonists (52.6%) and tryptamines (21%). The *in vitro* model most used was cultured cortical neurons extracted from rodents at different stages of embryonic development (26%). Risk of bias quality assessment was not performed for *in vitro* reports.

#### In vivo studies

In our sample, 51 original articles reported *in vivo* experiments in animal models. The most commonly used psychedelics administered were cannabinoid agonists (29.4%), NMDA antagonists (27.4%), and tryptamines (17.6%); however, a wide range of drugs and dosages were used in these studies. The animal models most present were rats (66.6%), followed by mice (31.4%). Other models, such as chick embryos, *Xenopus* frogs, and *Drosophila* larvae, were present in one study each.

We performed a quality of report analysis using the SYRCLE Risk of Bias tool (Hooijmans et al., 2014), and results are summarized in **Figure 2**. Nearly all *in vivo* studies did not report a valid method of randomization, so we reduced the quality level of the report as recommended by the SYRCLE initiative. We graded these studies as “Yes” if the authors mentioned any method of randomization, regardless of its robustness. The second major source of bias in these studies was the lack of control over housing conditions, as only one article described the housing of experimental animals during the experiment in a satisfactory manner. In addition, for criteria 7 (40%), 3 (24%), 5 (14%), and 6 (12%), the number of studies graded positively did not reach 50% of the total pool, indicating a lack of rigor in most *in vivo* studies, which is a common issue not only in psychedelic science but across a range of experimental and methodological articles within the life sciences (Carneiro, Moulin, Macleod, & Amaral, 2018; Macleod et al., 2015; Moulin, Rayêe, Williams, & Schiöth, 2020). These results highlight the importance of improving quality of reporting for increasing reproducibility in biological research (Begley & Ioannidis, 2015; Ioannidis, 2014). In the following sections, we will delve into each chemical category in more depth, with percentages in parentheses indicating the proportion of the total number of articles (n=68) unless otherwise stated.

### Summary of findings

#### CB1 agonists

Cannabinoids are unique in the central nervous system (CNS). The CB1 receptor was first cloned in Tom Bonner’s lab (Matsuda et al., 1990) and is predominantly expressed in neurons, where it is responsible for the psychoactive effects of cannabinoids (Raphael Mechoulam & Parker, 2013). The CB1 receptor remained an “endogenous orphan” until 1999, when Raphael Mechoulam’s team discovered the arachidonoylethanolamide (AEA) and the 2-arachidonoylglycerol (2-AG)(Martin, Mechoulam, & Razdan, 1999). In contrast, the CB2 receptor is less prevalent in the brain but is widely expressed in the immune system (Onaivi, 2006; Pertwee, 2006). While we have a good understanding of the effects of cannabinoids on humans based on observational studies of subjects who smoke plants in the *Cannabis* genus, which contain the phytocannabinoids THC and CBD (Bonini et al., 2018; Chang & Chronicle, 2007), it was only recently that mechanisms underlying their homeostatic function were first understood (de Fonseca et al., 2005).

The endocannabinoids system (eCB system) is a complex and widespread system that acts retrogradely as a feedback channel, influencing the release of all classical neurotransmitters (Skaper & Di Marzo, 2012) and enabling short- and long-term adaptive responses that lead to plasticity changes through mechanisms that are not yet fully understood (Araque, Castillo, Manzoni, & Tonini, 2017). While not all cannabinoids are psychoactive and only some are considered psychedelics (Bonini et al., 2018), we have chosen to include CB1 agonists because their availability as medicine is increasing worldwide, and because their synthetic analogs have varying potencies and behavioral effects that have not yet been fully described.

In our review, we identified 21 *in vitro* and *in vivo* studies examining the effects of five different cannabinoids: synthetic compounds HU210 (23.8%), Win55212-2 (9.5%), and O-2545 (4.7%), the endocannabinoid AEA (14.2%), and the phytocannabinoid THC (61.9%). Only one report included humans, a case-control study (Demirakca et al., 2011). Most of the experiments were conducted *in vivo* (71.42%), with a focus on adolescent murine models (47%) in an effort to better understand the widespread misuse of Cannabis plants, which is the most commonly used illicit drug globally, particularly among youth (Peacock et al., 2018). All of the studies used molecular or cellular methods of measurement, with a lesser focus on behavioral (38%) and electrophysiological (9.5%) measures. Descriptive results are summarized in **Supp Table 1**.

The relationship between neurogenesis and THC, as detailed in **Table 1**, appears conflicting due to differing study designs. For instance, the THC dosage varies greatly among studies, with some administering significantly high doses up to 30 mg/kg (Downer, Gowran, & Campbell, 2007; Kochman, dos Santos, Fornal, & Jacobs, 2006) — levels not generally therapeutic and potentially excessive even for high tolerance individuals (Karila et al., 2014). Conversely, other studies have examined a wide range of THC doses, from 0.75 to 10 mg/kg (Beiersdorf et al., 2020; Cuccurazzu et al., 2018; Downer et al., 2007; Kochman et al., 2006; Leishman, Murphy, Mackie, & Bradshaw, 2018; Poulia et al., 2021; Realini et al., 2011; Rubino et al., 2008; Steel, Miller, Sim, & Day, 2014; Suliman, Taib, Moklas, & Basir, 2018). The time point of intervention and analysis also significantly diverges across the studies. Three distinct intervention periods are identifiable: embryonic/early post-natal phase exposure (Beiersdorf et al., 2020; Berghuis et al., 2007; Downer et al., 2007; Psychoyos, Hungund, Cooper, & Finnell, 2008); exposure during adolescence, specifically in rats between P28-P45 (Cuccurazzu et al., 2018; Lee et al., 2014; Poulia et al., 2021; Rubino et al., 2008; Steel et al., 2014; Suliman et al., 2018); and exposure during adulthood (Jiang et al., 2005; Kochman et al., 2006; Leishman et al., 2018; Rueda, Navarro, Martínez-Serrano, Guzmán, & Galve-Roperh, 2002), as well as in-vitro studies with various models (Shum et al., 2020; Stanslowsky et al., 2017; F. Zhang, Challapalli, & Smith, 2009; Zhou & Song, 2001).

As shown in **Table 1**, several studies reported a negligible impact of acute or chronic CB1 agonists on neurogenesis (Berghuis et al., 2007; Downer et al., 2007; Kochman et al., 2006; Stanslowsky et al., 2017; Steel et al., 2014). Importantly, it should be noted that some studies focused not directly on neurogenesis but on other indirect plasticity markers (Berghuis et al., 2007; Poulia et al., 2021; Stanslowsky et al., 2017) or teratogenic morphological effects (Beiersdorf et al., 2020; Downer et al., 2007; Psychoyos et al., 2008). Yet, three studies identified positive effects on adult plasticity, specifically regarding the survival of newborn neurons and neurite growth, both *in vivo* and in culture (Jiang et al., 2005; Suliman et al., 2018; F. Zhang et al., 2009).

Further into our analysis, we identified notable findings that warrant a more detailed discussion, each contributing unique insights into the complex effects of cannabinoids on neural functioning and development. Parolaro’s group, for instance, employed a chronic, escalating THC regimen (11 days, twice a day) spanning three distinct age windows (2.5mg/kg in P35-P37; 5mg/kg in P38-P41; 10mg/kg in P42-P45). Their findings highlighted three significant outcomes: Firstly, they found that chronic THC exposure during adolescence indirectly reduced Brain-Derived Neurotrophic Factor (BDNF) via the CREB pathway in the hippocampus and prefrontal cortex (PFC) of males, while paradoxically increasing BDNF in the nucleus accumbens (NAc) of females (Rubino et al., 2008). Secondly, they observed a reduction in cell proliferation in females, an effect that was prevented by the fatty acid amide hydrolase (FAHH) inhibitor URB597 (Poulia et al., 2021; Realini et al., 2011) Finally, extending their earlier work, they demonstrated that such a THC exposure regimen could decrease not only the proliferation but also the survival rates of adult-born granule cells (abGC) in female rats (Cuccurazzu et al., 2018).

Broadening the perspective, Antoniou’s group in Greece demonstrated that chronic exposure to THC during adolescence disrupts the balance of dopamine transmission in several brain regions, including the prefrontal cortex, hippocampus, nucleus accumbens, and striatum. These findings provide a broader understanding of the molecular consequences of cannabis abuse during adolescence (Poulia et al., 2021). Lee et al. used a similar chronic exposure design, escalating through adolescence with the synthetic cannabinoid HU210, with dose ranges in the same age windows mentioned earlier (0.025 – 0.050 – 0.100 mg/kg). Their work resulted in the discovery of reduced proliferation and survival rates, but only in male mice evaluated at P70 (Lee et al., 2014).

In a study involving a transgenic model of Alzheimer’s Disease (AD) APP23/PS45, Chen et al. administered a daily dose of HU210 (0.010mg/kg) chronically for 10 and 20 days to both old and adolescent mice. Their results showed a reduction in BrdU+ cell numbers in young mice, and a notable observation was that most of the animals (16/20) did not survive until the end of the planned treatment (20 days), which led the authors to halve the treatment duration, indicating the potentially harmful effects of HU210, especially given its 100-800 times higher potency compared to THC (Chen et al., 2010; R. Mechoulam et al., 1988). However, the authors’ focus was primarily on AD-related features, which limits the extrapolation of these findings to a scenario involving lower doses and healthy conditions.

Leishman and colleagues took a different approach, administering a single 3 mg/kg THC injection to female CD1 rats at three different age windows (P35 – P60 – P90), followed by lipidome and transcriptome analyses 2 hours later. They found eCB levels were reduced across the brain, with the greatest impact observed in the adult hippocampus (Leishman et al., 2018). Additionally, Beiersdorf et al.’s study involved proteomics analyses of a pre-natal intervention (THC 1, 5 mg/kg from P5-P16 or P5-P35 days s.c.) and reported severe deficits in mitochondrial function at P48 and P120, indicating long-term consequences from early THC exposure (Beiersdorf et al., 2020).

Moreover, Rueda et al. reported that 24-hour *in vitro* exposure to the eCB AEA (5 µm) reduced neurite growth, with this reduction mediated by ERK signalling. Their findings were based on a variety of biochemical tools, and they also noted a decrease in the survival rates for abGC in adult rats (exact age not reported) following a 4-day *in vivo* exposure to M-AEA (daily 5mg/kg IP). (Rueda et al., 2002). Similarly, Zhou and Song’s study, which involved acute exposure of neuroblastoma N1E-115 cells to HU210 (0.01-100nM), showed that neurite growth was greatly dampened in a dose-dependent manner. This effect was mediated by intracellular cAMP (Zhou & Song, 2001). Lastly, Shum et al. proposed another mechanism for CB1R-mediated dampening of neurite growth in vitro: the involvement of ERK and Akt signalling pathways. They showed that the neurite growth ratio could be restored to control levels by the CB1R antagonist SR141716, at least in the case of human-induced Pluripotent Stem Cells (hiNSC) differentiated into neurons (Shum et al., 2020).

We believe that extrapolating these results to a wider population warrants caution, particularly considering the gender discrepancies observed in murine models and their possible lack of translationability to humans. Such differences challenge the direct applicability of these findings to patients (Naqvi et al., 2019). Legalization of cannabis in certain regions provides a unique opportunity for researchers to study its gender-specific effects and side effects in human populations. This is illustrated by the findings of two Canadian studies (Matheson et al., 2020; Matteau-Pelletier, Bélanger, Leatherdale, Desbiens, & Haddad, 2020).

From a neurobiological standpoint, it is postulated that CB1 agonists could potentially influence neurogenesis by modulating 5-HT and ACh neurons in the DRN and MS, respectively. These neurotransmitters are known to positively regulate different steps in neurogenesis (Raphael Mechoulam & Parker, 2013). However, the intricate relationship between the eCB system and other neurotransmitters, further complicated by species-specific variations, adds a layer of complexity to this field of study. Amidst these persisting ambiguities surrounding the mechanistic action of cannabinoids on the brain, it is pertinent to note the reported benefits of both phytocannabinoids and their synthetic counterparts. These compounds have been observed to confer antidepressant and anxiolytic effects in patients experiencing depression and those undergoing chemotherapy (Rodriguez Bambico, Duranti, Tontini, Tarzia, & Gobbi, 2009). This is in the backdrop of a longstanding relationship between our species and these compounds, a relationship that can be traced back to as early as 12,000 years A.C (Ren et al., 2021), In light of their proven safety, it is prudent to continue the exploration of the potential benefits these compounds may offer. Importantly, considerations of the implications of their use and misuse should be informed by a global perspective, taking into account data collected from human populations across the world (Ferber et al., 2019; Garani, Watts, & Mizrahi, 2021)

#### NMDA antagonists

Ketamine and phencyclidine (PCP), synthetic drugs originally discovered at the Parke-Davis Research Center, are arguably the most studied NMDA antagonists (Hoefle & Warner-Lambert, 2001; Pochwat, Pałucha-Poniewiera, Szewczyk, Pilc, & Nowak, 2014). Initially employed as anaesthetics in the late 20th century (Domino & Warner, 2010; Journey & Bentley, 2021), the compound induces a dissociative state (Domino & Warner, 2010; Journey & Bentley, 2021), which may lead to side effects ranging from hallucinations to mania (Javitt & Zukin, 1991; Pochwat et al., 2014; Ruiz & Strain, 2011). The discovery of ketamine’s rapid antidepressant effect (Berman et al., 2000) was considered by some researchers the greatest breakthrough in depression research over the past half-century (Data-Franco et al., 2017; Hashimoto, 2019; Sial, Parise, Parise, Gnecco, & Bolaños-Guzmán, 2020). Within our review, 20 of the included studies utilized NMDA antagonists, accounting for the second most studied chemical group of psychedelics. The time point of analysis differs between laboratories, but the single-dose paradigm, employed by 80% of ketamine studies, remains the most remarkable aspect of ketamine treatment (see descriptive results in **Supp. Table 2**). Ketamine’s facilitatory effect on plasticity has been thoroughly examined through physiological and behavioural measures and, to a lesser degree, electrophysiological methods (**Table 2**). Across diverse protocols, the focus was primarily on elucidating the cellular pathway through which ketamine boosts Brain-Derived Neurotrophic Factor (BDNF), subsequently promoting synaptogenesis, neurite growth, and neurogenesis (de Vos, Mason, & Kuypers, 2021). Among the *in vivo* ketamine studies (n = 10), the most frequently used behavioural assessment was the forced swim test (FST; 40%), followed by the open field test (OF; 20%). Other tests, such as the novel object recognition test (NOR), novelty suppressed feeding test (NSFT), splashing test (ST), and Morris water maze (MWM), were each utilized in 10% of the studies. Broadly, these behavioural tests were deployed to evaluate cognitive abilities known to be impaired in major depressive disorders.

Ketamine has been shown to enhance synaptogenesis and neurite growth in young adult mice when administered at a subanaesthetic dose ranging from 0.25-10 mg/kg/day (Akinfiresoye & Tizabi, 2013; N. Li et al., 2010; R.-J. Liu et al., 2013; Ly et al., 2018). However, three studies using doses beyond this range reported potentially harmful effects on neurogenesis and associated behaviours. First, Zuo and colleagues administered 30mg/kg/day of ketamine to male rats between postnatal days 35-42 (P35-P42), along with 2-4g/kg/day of alcohol over 14 days. They observed a reduction in synapsin expression in the hippocampus, a phenomenon that occurred even in the absence of alcohol. The study also reported that ketamine treatment in adolescent mice induced hyperlocomotion, anxiety (assessed using OF), ataxia, stereotyped behaviour, mitochondrial damage, and morphological abnormalities in hippocampal and cortical cells. Additionally, dopamine (DA) and glutamate levels increased in the cortex and hippocampus, suggesting a possible disruption of the CREB pathway, causing the aforementioned adverse effects (Zuo et al., 2018). Next, Huang et al. (2016) delivered four sub-anesthetic intraperitoneal (IP) injections of 10mg/kg ketamine over 4 hours to P7 rats. This regimen led to a reduction in neuronal and glial proliferation without impacting neuronal differentiation (as assessed by B-Tubulin III/BrdU+ co-labelling) up to 14 days post-injection. However, the labelled cell levels in all assays returned to control levels 21 days post-injection (Huang et al., 2016). Lastly, Schiavone et al. (2020) elucidated the mechanisms underlying the neurochemical imbalance and oxidative stress observed in adulthood following intermittent administration of 30 mg/kg ketamine to neonatal rats (P7, P9, P11). A two-fold increase in NOX2 immediately preceding the chemical insult persisted into adulthood, resulting in other observed imbalances. Although the authors did not measure the number of neural progenitors in the SGZ, it’s been found that NOX2 knockout mice exhibit a reduced number of radial glial-like (RGL) cells compared to wild-type mice in adulthood (Dickinson, Peltier, Stone, Schaffer, & Chang, 2011), indicating a possible bimodal, dose-dependent effect of ketamine on RGL proliferation during this specific ontogenetic window of exposure via NOX2 overexpression (Schiavone et al., 2020).

The prevailing consensus in the scientific literature suggests that ketamine positively influences the subprocesses of adult neurogenesis via the upregulation of BDNF (Browne & Lucki, 2013; Duncan et al., 2013; Haile et al., 2014; Keilhoff, Bernstein, Becker, Grecksch, & Wolf, 2004; Lepack, Fuchikami, Dwyer, Banasr, & Duman, 2015; Rybakowski et al., 2013). However, a study by Goulart et al. (2010) presents an intriguing contrast. They observed a dose-dependent reduction in the BDNF increase stimulated by the Novel Object Recognition (NOR) task, assessed approximately 5 hours post-training/injections (Goulart et al., 2010). This seemingly contradictory finding may be partially explicated by the research conducted by Huang and colleagues. Although they did not measure neurotrophin levels, their observations suggested a transient reduction in proliferation and differentiation, possibly mediated by BDNF (Huang et al., 2016). This could provide a plausible mechanism for the ketamine-induced disruption of the consolidation phase of long-term recognition memory observed by Goulart and colleagues. Interestingly, despite the disruption in the migration of adult-born Granule Cells (abGC), leading to an increased number of GCs in the SGZ until at least P44, the cognitive deficits persisted until at least P60 as assessed by Morris Water Maze (MWM) performance (Huang et al., 2016). Nevertheless, it is important to note that Huang et al.’s study involved a potent intoxication early in life (P7), and their results do not mirror those obtained in adult animals (Browne & Lucki, 2013; Data-Franco et al., 2017; Duman & Li, 2012; Duncan et al., 2013; Ly et al., 2018; Olson, 2018). This discrepancy reinforces the need for further research to elucidate the dynamic interactions between ketamine and neurotrophins under conditions that stimulate neurogenesis.

The majority of studies on phencyclidine (PCP) have sought to elucidate the neurocognitive damage caused by chronic exposure in utero or neonatally. All of these studies employed physiological assessments, and nearly half of them incorporated tests for behavioural evaluation (Tanimura et al., 2009; Toriumi et al., 2012). The Bromodeoxyuridine (BrdU) label for cells undergoing the S-phase was the most frequently used method (80%). The consensus from these studies is that early exposure to PCP has a significantly detrimental impact on neurogenesis (Maeda et al., 2007; Tanimura et al., 2009; Toriumi et al., 2012) and overall plasticity (Toriumi et al., 2012; Q. Zhang, Yu, & Huang, 2016).

In contrast, a study by Liu et al. (2006) in Japan reported different results. They administered 7.5 mg/kg PCP acutely and chronically (for 5 or 14 days) for comparison within groups. They found that acute and 5-day chronic exposure resulted in minor or no effects on cell proliferation, differentiation, and survival in the dentate gyrus of adult rats when compared to the saline control. Interestingly, the decrease induced by these treatments was restored to control levels after one week of abstinence (J. Liu et al., 2006).

However, their study lacks detailed information on the exact age of the animals, a factor that strongly influences adult neurogenesis (Kuhn, Dickinson-Anson, & Gage, 1996), only stating that it was conducted on young adult animals, not neonatal or in utero subjects. In summary, it is reasonable to assert that exposure to these psychoactive substances during the early stages of neural development can induce detrimental effects. Moreover, the safety of PCP as a therapeutic agent needs further evaluation in adult models, preferably in disease-relevant models, rather than focusing solely on its teratogenic effects.

Human trials with ketamine shed light on various processes contributing to plasticity. For instance, a cross-sectional study conducted by Fan and colleagues revealed that ketamine addiction can lead to low-grade inflammation (Fan et al., 2015). Further, three other studies specifically focused on patients with major depressive disorder (MDD), particularly treatment-resistant depression (TRD) (Duncan et al., 2013; Haile et al., 2014; Rybakowski et al., 2013). These studies found a correlation between non-responsiveness to ketamine treatment in TRD patients and decreased BDNF levels 7 and 14 days after a single 0.5mg/kg ketamine infusion (Rybakowski et al., 2013). Interestingly, when applied in a similar protocol, this same dose led to an increase in serum BDNF levels as assessed 230 minutes after the infusion. This increase was accompanied by an increase in Slow Wave Sleep (SWS) duration in responders (individuals characterized by a decrease of more than 50% in their baseline Montgomery-Asberg Depression Rating Scale (MADRS) score (Duncan et al., 2013). A similar study conducted in Texas, USA, yielded comparable results (Haile et al., 2014).

Moreover, in a separate human-controlled trial at the New York State Psychiatric Institute, ketamine administered to cocaine-addicted patients who weren’t seeking treatment or abstinence, reduced their cocaine self-consumption and had high scores on the Hood Mysticism Scale (HMS) when compared to other scales like the Clinician-Administered Dissociative States Scale (CADSS) and the Near-Death Experience Scale (NDES) (Dakwar et al., 2018). These results provide some insight into which subjective effects of the ketamine psychedelic experience might be beneficial for addiction treatment, suggesting these aspects should be monitored in assisted psychotherapy.

### Harmala alkaloids

Harmala alkaloids are indole nitrogenated compounds featuring a β-carboline heterocyclic structure, first identified in the seeds of *Peganum harmala*, also known as Syrian Rue (Herraiz, González, Ancín-Azpilicueta, Arán, & Guillén, 2010). Recently, these compounds have attracted increased attention due to their presence in *ayahuasca* brews. The primary compounds in this category include harman, harmine, harmaline, and tetrahydroharmine (de Oliveira Silveira et al., 2020). While these β-carbolines are not inherently psychedelic, they can influence serotonin signaling by inhibiting the monoamine oxidase enzyme (MAO), thus producing behavioral effects somewhat similar to Selective Serotonin Reuptake Inhibitors (SSRIs) (Domínguez-Clavé et al., 2016). Often referred to as MAO inhibitors (MAOis), these compounds are ubiquitous in the various forms of *ayahuasca* prepared by indigenous populations (McKenna, Towers, & Abbott, 1984). Interestingly, in some versions of *ayahuasca*, DMT is even absent (Callaway, 2005; Rivier & Lindgren, 1972). Their antidepressant effect has been explored in psychiatry, with some MAOis even being marketed under names like Neuralex and Marplan, among other synthetic functional analogs (Fagervall & Ross, 1986; Robinson, 1973). However, most MAOis were withdrawn from the market due to hepatotoxicity (Lopez-Munoz & Alamo, 2009), although it may not be the case of the harmala alkaloids (Brito-da-Costa, Dias-da-Silva, Gomes, Dinis-Oliveira, & Madureira-Carvalho, 2020). The cellular mechanisms of such compounds are still being explored, more recently under the light of the neurogenic hypothesis for MDD, as their effects on neurogenesis and plasticity processes are uncovered (Ferraz, de Oliveira Júnior, Picot, da Silva Almeida, & Nunes, 2019).

All studies involving harmala alkaloids (6 included, **Supp. Table 3**) focus on their potential antidepressant effects. *In vivo* studies also assessed hippocampal BDNF in conjunction with behavioral methods. Half of articles from this category originated from a Brazilian research team led by João Quevedo, with Jucélia Fortunato as a key contributor, which conducted a series of rat experiments revealing the potential antidepressant effects of the harmine alkaloid (Fortunato et al., 2010a, 2010b, 2009). The team initially found that male rats at post-natal day 60 (P60) showed improved performance in the forced swim test (a measure of learned helplessness, a common depressive-like behavior) after being treated with 10 or 15mg/kg of harmine one hour prior to the test. Their locomotor activity (measured by open field test) remained normal. The improvement was comparable to that produced by the tricyclic antidepressant imipramine. Intriguingly, rats treated with 15 mg/kg of harmine demonstrated a two-fold increase in BDNF levels immediately after the test, while those treated with imipramine did not (Fortunato et al., 2009). The following year, the team reported similar results after chronic administration (14 days) of harmine (Fortunato et al., 2010a).

A critique of their initial studies pertained to the relevance of the results to depression pathophysiology. To address this, they later generated a study incorporating a Chronic Mild Stress (CMS) model of depression in rats. There, they administered 15 mg/kg of harmine over a 7-day treatment period, using a similar protocol but without the imipramine control. Similar results were found. Moreover, the forced swim test (FST) assessment was replaced with the sucrose preference test (SPT) to measure anhedonia-like phenotypes, a core symptom of depression. The study demonstrated that harmine-treated mice exhibited less anhedonia only after CMS. The study also found that harmine treatment reduced blood levels of adrenocorticotropin (ACTH), a stress hormone, while adrenal weight was similar to that of non-CMS rats. Contrary to their previous findings, BDNF levels after treatment were not affected. Such discrepancy could be due to the different time-points of sample collection employed in the study (first after a stressful situation, second after a mildly pleasant situation), suggesting that harmine may dynamically influence BDNF levels (Fortunato et al., 2010b).

Harmine was also studied in Jiangsu, China, by a team led by Professor Chao Huang and Liu and colleagues, who used Chronic Unpredictable Stress (CUS) to model depressive symptoms in mice. They examined the effects of chronic administration of 10 and 20 mg/kg of harmine using the forced swim test (FST), sucrose preference test (SPT), and tail suspension test (TST) — the three most commonly used tests for measuring depressive symptoms in mouse models. It was reported a 40% effect favoring the antidepressant action of harmine in all behavioral tests, which was comparable to the positive control, fluoxetine (F. Liu et al., 2017). In addition to behavioral tests, biochemical markers were investigated. The team described that 20 mg/kg of harmine increased BDNF levels in the hippocampus and prefrontal cortex, along with glutamate transporter 1 (GLT-1) expression levels. They also found that harmine promoted the survival of newborn cells, as evidenced by doublecortin (DCX) immunofluorescence, and prevented the reduction in glial fibrillary acidic protein (GFAP) induced by the CUS protocol (F. Liu et al., 2017). Lastly, the authors tested the hypothesis that these regulatory effects might be due to the restoration of astrocytic function by administering L-Alpha-Aminoadipic Acid (L-AAA), an astrocyte-specific gliotoxin, intracerebroventricularly (i.c.v). Indeed, this treatment shifted the results towards those of the saline-CUS group (F. Liu et al., 2017).

Two other studies conducted entirely in vitro reported that harmine increased cell proliferation. One study from Rio de Janeiro, Brazil, showed that the presence of 7.5 µM harmine in the culture medium of human neural progenitor cells (hNPCs) increased proliferation by 71.5% (Dakic, de Moraes Maciel, et al., 2016). A Spanish team went further and tested four different harmala alkaloids (harman, harmine, tetrahydroharmine, and harmol) at a 1 µM concentration, acutely (3 days) and chronically (7 days), on neurospheres derived from the subgranular zone (SGZ) and subventricular zone (SVZ) of adult mice. The treatment enhanced all stages related to neurogenesis in vitro: proliferation, migration, and differentiation (Morales-García et al., 2017a). Altogether, the current body of research suggests that harmala alkaloids have a positive impact on neurogenesis and neuroplasticity, particularly within the framework of depression models (**Table 3**).

The antidepressant potential of MAO inhibitors (MAOis) is well known and has been explored in the past. However, more research is needed regarding the side effects of harmala alkaloids (Wimbiscus, Kostenko, & Malone, 2010). Future studies should focus on expanding our understanding of the long-term safety of these compounds, comparing them with synthetic MAOis, preferably those still on the market. This research will help determine whether their positive effects can counterbalance potential hepatotoxicity and severe interactions with tyramine-rich foods. By identifying which part of the harmala alkaloid structure is responsible for differences in action and consequences, we can pave the way for the development of safer analogs.

### Psychoactive tryptamines

Tryptamines are indolamines neurotransmitters that originate from tryptophan (Araújo, Carvalho, Bastos, Guedes de Pinho, & Carvalho, 2015). All psychoactive tryptamines are potent 5-HT2A agonists, and many also demonstrate affinity for 5-HT1 and other 5-HT2 receptors. Notably, they may also interact with ionotropic and metabotropic glutamate receptors, dopamine, acetylcholine, and trace amine-associated receptor (TAAR) (Carbonaro & Gatch, 2016). Certain DMT analogs have shown activity via sigma-1 receptors as well (Koornneef et al., 2009). This broad receptor affinity equips psychoactive tryptamines with the capability to induce altered states of consciousness, characterized by heightened introspection (de Araujo et al., 2012) and changes in sensory perception, mood, and thought (Calvey & Howells, 2018).

While psychoactive tryptamines, used by ancient civilizations globally for millennia (Desmarchelier, Gurni, Ciccia, & Giulietti, 1996), were first clinically investigated in the mid-20th century for the treatment of mood and drug abuse disorders, and some advocates proposed their systematic use by healthy individuals to enhance cognition (Nichols, 2016; Williams, Rao, & Goldman-Rakic, 2002). Despite their shared primary pharmacologic pathway—5-HT2A receptor agonism—these tryptamines can generate different experiences due to their variable affinity for other receptors. The current review includes eleven studies that meet the inclusion criteria, investigating the effects of psilocybin (18%), ibogaine (9%), and DMT analogs, including the ayahuasca concoction (73%). These tryptamines were predominantly assessed using physiological and behavioral methodologies, with electrophysiological methods used less frequently (**Supp. Table 4**).

As shown in **Table 4**, the majority of the included studies provide evidence that, irrespective of their specific type, tryptamines have the potential to enhance aspects of neuroplasticity (Catlow, Song, Paredes, Kirstein, & Sanchez-Ramos, 2013; Dunlap et al., 2020; Katchborian-Neto et al., 2020; Lima da Cruz et al., 2018; Ly et al., 2018; Marton et al., 2019; Morales-Garcia et al., 2020). Remaining studies, although not directly assessing neurogenesis, reported no adverse effects on behavior or cellular physiology (Castro-Neto et al., 2013; Wankhar, Syiem, Pakyntein, Thabah, & Sunn, 2020; Winne et al., 2020). For instance, Catlow et al. (2013) discovered that a single low dose (0.1-0.5 mg/kg) of psilocybin, administered 24 hours prior to a Trace Fear Conditioning test, initially heightened fear response but facilitated fear extinction by the third trial, an effect not observed until the tenth trial in control animals. Conversely, high doses (1-1.5 mg/kg) showed no behavioral impact and reduced BrdU-labeled cells 18 days post-treatment. (Catlow et al., 2013).

Another team, based in Shillong, India, investigated the acute and sub-acute (7 days) effects of 0.7 mg/kg 1-Methylpsilocin, a psilocin tryptamine derivative. Their research aimed to assess the potential of 5-HT2CR exploitation in mitigating symptoms common to mood disorders. Following a Chronic Unpredictable Stress (CUS) protocol in rats, they employed Open Field (OF), Forced Swim Test (FST), and Elevated Plus Maze (EPM) tests in conjunction with tissue sample analysis. Their findings indicated that 1-Methylpsilocin alleviated behavioral signs of anxiety and anhedonic phenotype by the seventh day post-CUS. Furthermore, they demonstrated that this treatment could rescue subjects from CUS-induced mitochondrial damage and imbalanced 5-HT/5-HT2CR levels in the PFC and hippocampus. Remarkably, the treatment also mitigated hippocampal cell morphological abnormalities and the CUS-induced rise in corticosterone levels (Wankhar et al., 2020).

Ibogaine, a naturally occurring psychoactive tryptamine found in certain plants of the Apocynaceae family native to Central Africa, has been investigated for its potential to mitigate drug-seeking behavior (Alper, Lotsof, & Kaplan, 2008). Researchers in Uruguay examined the compound’s influence on neuroplasticity in mesocorticolimbic areas of the rat brain, uncovering intricate interactions between NGF, GDNF, and BDNF, both in terms of mRNA and protein products. Briefly, 20 or 40 mg/kg doses of ibogaine (I_20_ and I_40_) dose-dependently upregulated BDNF and NGF mRNA in PFC, Ventral Tegmental Area (VTA), Substantia Nigra (SN), and Nucleus Accumbens (Nacc) 24 hours post-injection. However, 3 hours post-injection, a decline of 1.7 and 2-fold in PFC-BDNF mRNA was observed for the I_20_ and I_40_ groups, respectively. Notably, I_40_ also elevated GDNF levels in VTA and SN 24 hours later. In terms of protein levels, upregulation was confined to proBDNF-NAcc (for both I_20_ and I_40_) and VTA-GDNF exclusively for I40, with a possible tendency for increased BDNF in NAcc and VTA that warrants further investigation (Marton et al., 2019).

The indole alkaloid 5-Methoxy-N, N-dimethyltryptamine (5-MeO-DMT), a structural analog of DMT found in the venom of the *Incilius alvarius* toad, has been the subject of several studies (Araújo et al., 2015). In a series of three experiments, our team investigated the effects of 5-MeO-DMT on adult neurogenesis (Lima da Cruz et al., 2018). Our first two experiments showed an increase in the proliferation and survival of newborn granule cells in the ventral dentate gyrus following a single intracerebroventricular injection of 100µg of 5-MeO-DMT. In the third experiment, we employed the whole cell patch clamp technique to measure the passive and active membrane properties of the identified doublecortin-positive cells. The results revealed that the adult-born granule cells (abGCs) from treated animals exhibited electrophysiological properties more akin to mature granule cells. In a separate study, Winne et al. examined salicylate-induced anxiety-like behavior. Their findings suggested that salicylate can induce theta-2 and slow gamma oscillations in young mice, correlating with anxiogenic behavior. This effect was mitigated by a single 20 mg/kg intraperitoneal injection of 5-MeO-DMT administered a week prior to the test (Winne et al., 2020).

Moreover, researchers from the University of São Paulo, Brazil, conducted a study on adult rats to assess changes in neurotransmitter levels 40 minutes post-ingestion of *ayahuasca*. Their findings indicated that all tested ayahuasca concentrations (250, 500, 800 mg/kg oral) elevated GABA and 5-HT levels in the hippocampus and reduced glycine and GABA levels in the amygdala, following a dose-dependent pattern. Moreover, levels of all monoamines in the amygdala increased, yet the rate of serotonin consumption was reduced in the amygdala. Only the highest dose of 800 mg/kg could reproduce this effect in the hippocampus (Castro-Neto et al., 2013). Regarding DMT analogs, noteworthy efforts are being made by David E. Olson’s lab at UC Davis to develop psychedelics without hallucinogenic properties while maintaining their neuroplastic effects, thus potentially reducing undesirable effects for some patients. Despite certain criticisms, Dunlap and colleagues reported successful results, demonstrating that isoDMT and 5-MeO-isoDMT enhanced neurite growth in cultured cortical neurons, much like their natural counterparts. Interestingly, in further experiments, these compounds did not elicit the head-twitch response (HTR) - a behavior commonly observed in mice when a hallucinogenic 5-HT2A agonist is administered (Dunlap et al., 2020).

### Entactogens/ empathogens

As atypical psychedelics (Calvey & Howells, 2018), those compounds typically induce an emotional sense of “oneness,” notably enhancing empathy on top of the usual psychedelic experience (Nichols, 1986). These unique properties distinguish them from other classes of drugs. All entactogens bind to 5-HT_1A_ and 5-HT_2_ receptors with varying affinities. Some phenethylamines, such as MDMA (3,4-methylenedioxymethamphetamine) and its analogs, affect other monoamines by blocking their uptake and forcing their release into the synaptic cleft (Papaseit et al., 2020).

Although some tryptamines are included as entactogens (Nagai, Nonaka, & Satoh Hisashi Kamimura, 2007) our review found only articles exploring the effects of MDMA within our scope. MDMA was initially developed in 1912 by Merck chemist Anton Köllisch to be an appetite suppressant, but it was never sold for this purpose (Kalant, 2001). In the 1970s, psychopharmacologist Alexander Shulgin synthesized it and, after self-experimentation, suggested its use as a medicine to aid patients undergoing therapy for PTSD and addiction disorders (Shulgin & Shulgin, 1991). In the 1980s, MDMA became a street drug, often sold illicitly as “molly” or “ecstasy” (NIDA, 2020). Today, MDMA is in the final stages of clinical research for approval as a psychotherapeutic tool to treat PTSD (Emerson, Ponté, Jerome, & Doblin, 2014). The included articles explore the effects of MDMA during embryonic and/or neonatal stages (57.14%), adolescence (28.57%), and adulthood (28.57%) in murine models, with some articles examining more than one ontogenetic window (**Supp. Table 5**).

The findings of these studies, as summarized in Table 5, illustrate that the effects of MDMA on neurogenesis can vary depending on the dosage and duration of administration. In a study by Hernández-Rabaza et al., researchers attempted to mimic the binge consumption of MDMA often seen in social settings such as raves. They administered a dose of 40mg/kg over two days to male rats (the age of the rats was not reported). This treatment did not affect the proliferation of new cells. However, when assessed 14 days after the treatment, there was a noticeable 40% decrease in new cells (BrdU+ cells) in the dentate gyrus compared to the control group. The researchers also examined potential abnormalities in the growth of new neurons but found no difference between the MDMA-treated and control groups. (Hernández-Rabaza et al., 2006). Moreover, a study by Catlow et al. investigated the effects of a longer, chronic administration of MDMA. Rats were given a lower dose of 5 mg/kg per day for 10 days. The researchers observed an initial increase in new cells (BrdU+ cells) immediately after the MDMA treatment, but a significant decrease (around 40-50%) was noted 14 days later. The survival of these new neurons, measured by co-labeling BrdU and NeuN, was also reduced by approximately 40-50%. (Catlow et al., 2010).

The results presented by Catlow (2010) and Hernández-Rabaza et al. (2006) prompt further investigation into potential differential effects of MDMA across developmental stages. García & García (2016) report that a 60mg/kg dose of MDMA (3×5mg/kg over 4 days) more significantly impacted proliferation in adult rats (P48-P58) compared to adolescent rats (P27-P37), as demonstrated by ki67 and BrdU labeling (García-Cabrerizo & García-Fuster, 2016). Given the unreported age of the rats in Hernández-Rabaza et al.’s (2006) study, a direct comparison with García & García’s findings is not recommended. However, the reported age range of rats in Catlow et al.’s study (P28-39) corresponds to adolescence/young adulthood, introducing an element of ambiguity.

Further, García & García’s study also revealed a positive correlation between mature BDNF (mBDNF) levels and numbers of BrdU+ cells (r=0.492), with a stronger correlation observed between 5-HT2CR expression and mBDNF (r = 0.713). This led the authors to posit that chronic MDMA exposure (60 mg/kg) negatively affects overall plasticity, with more pronounced effects in adult animals. Interestingly, no significant impact was found when a lower dosage of 15mg/kg (3×5 mg within 1 day) was administered (García-Cabrerizo & García-Fuster, 2016).

With the growing popularity and widespread use of MDMA among young people and adults (Armenian & Rodda, 2020), three other studies focused on examining the effects of MDMA during early stages of life, given concerns about the drug’s potential teratogenic properties. In particular, a team of researchers from Seoul, Korea, investigated the impact on adult neurogenesis in embryos from dams exposed to 1.25 or 20 mg/kg MDMA for 36 days (E6-P21) via oral gavage. At 11 weeks post-exposure, animals were treated with BrdU. The study found that both doses led to reduced cell proliferation compared to saline, but only the higher dose of 20 mg/kg resulted in decreased survival of abGC as indicated by BrdU/NeuN labeling. Notably, only comparisons with female offspring in the experimental groups were reported in survival assays (Cho, Rhee, Kwack, Chung, & Kim, 2008).

In a teratology study conducted in Ohio, USA, Thompson et al. (2012) found that maternal ingestion of MDMA at a similar dose during E14-E20 increased fiber density in the prelimbic cortex (PLC) and CA1. Meanwhile, the expression of the norepinephrine transporter (NET) was elevated in the cornus ammonis regions, but not in the dentate gyrus (DG), Locus Coeruleus (LC), or PLC (V. B. B. Thompson et al., 2012). The authors suggested further exploration of NET upregulation to strengthen the association between downstream morphological (increased plasticity) and neurochemical abnormalities (upregulation of NE in the PFC and Nac) with previously reported cognitive and behavioral changes related to novelty (Koprich et al., 2003; V. B. B. Thompson et al., 2012; V. B. Thompson et al., 2009) However, the authors emphasized that the existing data does not suffice to determine causality. Subsequently, Canales and Ferrer-Donato (2014) demonstrated that a dose of 10mg/kg MDMA administered during a shorter pregnancy window (3 days, E13-E15) combined with alcohol significantly reduced neurogenesis (both proliferation and survival) by approximately 40% in adult female offspring compared to saline. Behavioral assessment via the Radial Arm Maze (RM) revealed about a 30% reduction in locomotion and a roughly 50% increase in working memory errors. Notably, the within-group variance was quite narrow, and only the group exposed to both drugs exhibited these impairments (Canales & Ferrer-Donato, 2014).

We conclude that abuse of MDMA with high acute doses results in enduring imbalances in the monoamine system and subsequent downregulation of neurogenesis. Conversely, low doses have no detectable effects on morphological abnormalities in adult brain tissue. It is noteworthy that the phase 3 studies currently being conducted by the Multidisciplinary Association for Psychedelic Studies (MAPS) are utilizing doses ranging from 0.08-0.18 grams of MDMA per patient (Mithoefer et al., 2019). Therefore, neurobiological studies should futher examine how low doses of MDMA impacts brain physiology.

### Other Psychedelics

In this section, we go through additional articles that employed psychedelic drugs from chemical categories not previously discussed. These include the substituted amphetamine with psychedelic effects 2,5-Dimethoxy-4-iodoamphetamine (DOI) (**Supp. Table 6**) (Belmer et al., 2018; Berthoux, Barre, Bockaert, Marin, & Bécamel, 2019; Jha, Rajendran, Fernandes, & Vaidya, 2008; Jones et al., 2009; Ly et al., 2018; Marinova et al., 2017) and the substituted phenethylamine 25I-NBOMe (**Supp. Table 5**) (Catlow et al., 2013).

A group from the French National Institute of Health and Medical Research (INSERM) assessed the capability of the 5-HT_2B_R to regulate Dorsal Raphe Nucleus (DRN) serotonin release. Activation of pet1+ DRN cells by the tryptamine BW723C86 (1µM), a 5-HT_2B_ receptor agonist, administered *ex vivo* increased the spontaneous firing rate. Intriguingly, in subsequent experiments, 5mg/kg DOI in vivo failed to boost the HTR in animals with a knockout of the 5-HT_2B_R, and 20 mg/kg MDMA in the same genotype also failed to replicate the increase in locomotor response observed in control mice. These behaviors were evaluated using the Open Field (OF) test (Belmer et al., 2018), indicating that 5-HT_2B_R may play a larger role in the behavioral effects of psychedelics on mice than previously believed.

A research team at the Tata Institute of Fundamental Research in Mumbai, India, using the BrdU labelling technique, reported that neither LSD (0.5 mg/kg) nor DOI (8 mg/kg) could increase proliferation in rats (age not reported) under either chronic (7 days) or acute treatment regimens (Jha et al., 2008). Another group from the Northwestern University Feinberg School of Medicine in Chicago, Illinois, reported that 1µM DOI could transiently promote dendritic remodeling in cultured cortical pyramidal neurons from rats, and this plasticity was mediated by Kalirin-7. The change involved the enlargement of spines, which began 30 minutes after exposure and returned to control size 60 minutes later (Jones et al., 2009). Ly and colleagues from UC Davis, in an article discussed previously (**Table 2**), extended the scope of their findings to other species, demonstrating that LSD and DOI could promote neuritogenesis in cultured neurons from mice and *Drosophila* larvae (Ly et al., 2018). Additionally, Marinova and colleagues employed human neuroblastoma SK-N-SH cells, TrkA is necessary for neurite extension (Marinova et al., 2017).

The only report found to investigate DOI using electrophysiological methods was conducted by Berthoux and colleagues. They used 1µM DOI on ex vivo slices from P14-P21 mice and reported that DOI induced Long-Term Depression (LTD) in mPFC layer 5 pyramidal neurons, as evidenced by whole-cell patch-clamp. The amplitude of the AMPA ESPC (excitatory postsynaptic current) coming from layer 1 Pyramidal stimuli was almost halved, and this decrease was mediated by 5-HT2AR since mice knocked out for 5-HT2AR did not show the same response. The internalization of AMPA receptors containing the GluA2 subunit through Protein kinase C (PKC) was necessary for DOI-induced LTD but not when LTD was replicated by a fast electrical pairing protocol (Berthoux et al., 2019). Moreover, in an article already discussed in the tryptamine section (**Table 2**), Catlow and colleagues used the substituted phenethylamine 25I-NBOMe in a single experiment to explore if 5-HT2AR activation affects proliferation of adult-born granule cells (abGC) in mice (age not reported). All tested doses (0.1, 0.3, and 1 mg/kg) reduced proliferation, but only the 1 mg/kg dose had a significant effect on the number of BrdU+/NeuN+ cells compared to saline-treated mice (Catlow et al., 2013).

In summary, these studies suggest that the substituted amphetamine DOI and the substituted phenethylamine 25I-NBOMe may have a range of effects on neural plasticity and behavior. These effects appear to be modulated by different serotonin receptors, including 5-HT2BR and 5-HT2AR. More research is needed to further understand the molecular mechanisms and the functional implications of these changes. Also, the specificity of the effects of these substances on different stages of neurogenesis needs to be further investigated.

## Conclusions

This systematic review sought to reconcile the diverse outcomes observed in studies investigating the impact of psychedelics on neurogenesis. Additionally, this review has integrated studies examining related aspects of neuroplasticity, such as neurotrophic factor regulation and synaptic remodelling, regardless of the specific brain regions investigated, in recognition of the potential transferability of these findings.

Our study revealed a notable variability in results, likely influenced by factors such as dosage, age, treatment regimen, and model choice. In particular, evidence from murine models highlights a complex relationship between these variables for CB1 agonists, where cannabinoids could enhance brain plasticity processes in various protocols, yet were potentially harmful and neurogenesis-impairing in others. These findings emphasize the need to assess misuse patterns in human populations as cannabinoid treatments gain popularity. We believe future researchers should aim to uncover the mechanisms that make pre-clinical research comparable to human data, ultimately developing a universal model that can be adapted to specific cases such as adolescent misuse or chronic adult treatment.

The only NMDA antagonist currently recognized as useful in medicine is ketamine. Many countries have adopted ketamine as a fast-acting antidepressant due to its remarkable effects on depressed individuals. However, research is still needed to evaluate its long-term effects on overall brain physiology. The studies discussed here have touched upon these issues, but further development is needed, particularly regarding the depressive phenotype, including subtypes of the disorder and potential drug interactions. For instance, recent findings indicate that ketamine and other psychedelics directly bind to the BDNF receptor TrkB (Tropomyosin receptor kinase B) (Moliner et al., 2023). Moreover, insights gained by psychiatrists from treating patients should generate ideas for pre-clinical research aimed at understanding ketamine’s interaction with neurotrophic factors.

Studies on harmala alkaloids, though limited, show promising antidepressant activity. This is not surprising, as this category of chemicals has already been explored in psychiatry. We believe that further research should focus on comparing harmala alkaloids with marketed analogues, particularly highlighting differences in side effects and common interaction problems. Moreover, incorporation of those substances by healthcare systems poses significant challenges. For instance, the ayahuasca brew, which combines harmala alkaloids with psychoactive tryptamines and is becoming more broadly studied, has intense and prolonged intoxication effects. Despite its effectiveness, as shown by many studies reviewed here, its long duration and common side effects deter many potential applications. Thus, future research into psychoactive tryptamines as therapeutic tools should prioritize modifying the structure of these molecules, refining administration methods, and understanding drug interactions. This can be approached through two main strategies: (1) eliminating hallucinogenic properties, as demonstrated by Olson and his team at UC Davis, who are developing psychotropic drugs that maintain mental health benefits while minimizing subjective effects; and (2) reducing the duration of the psychedelic experience to enhance treatment readiness, lower costs, and increase patient accessibility. These strategies would enable the use of tryptamines without requiring patients to be under the supervision of healthcare professionals during the active period of the drug’s effects.

Syncretic practices in Brazil, along with others globally, are exploring intriguing treatment routes using these compounds (Labate and Cavnar, 2014; Svobodny, 2014). These groups administer the drugs in traditional contexts that integrate Amerindian rituals, Christianity, and (pseudo)scientific principles. Despite their obvious limitations, these settings provide valuable insights into the drug’s effects on individuals from diverse backgrounds, serving as a prototype for psychedelic-assisted psychotherapy. In this context, it is believed that the hallucinogenic properties of the drugs are not only beneficial but also necessary to help individuals confront their traumas and behaviors, reshaping their consciousness with the support of experienced staff. Notably, this approach has been strongly criticized due to a rise in fatal accidents (Hearn, 2013; Holman, 2010), as practitioners are increasingly unprepared to handle the mental health issues of individuals seeking their services.

Nevertheless, this review highlights the potential benefits of psychedelics in terms of brain plasticity. Therapeutic dosages, whether administered acutely or chronically, have been shown to stimulate neurotrophic factor production, proliferation and survival of adult-born granule cells, and neuritogenesis. While the precise mechanisms underlying these effects remain to be fully elucidated, overwhelming evidence show the capacity of psychedelics to induce neuroplastic changes. Thus, rigorous preclinical and clinical trials are necessary to fully elucidate the mechanisms of action, determine optimal dosages and treatment regimens, and assess potential risks and side effects. It is crucial to investigate the effects of these substances across different life stages, from embryonic development to adulthood. Additionally,

## Past and Future Perspectives

Research with hallucinogens began in the 1960s when leading psychiatrists observed therapeutic potential in the compounds today referred to as psychedelics (Osmond, 1957; Vollenweider and Kometer, 2010). These psychotomimetic drugs were often, but not exclusively, serotoninergic agents (Belouin and Henningfield, 2018; Sartori and Singewald, 2019a) and were central to the anti-war mentality in the “hippie movement”. This social movement brought much attention to the popular usage of these compounds, leading to the 1971 UN convention of psychotropic substances that classified psychedelics as class A drugs, enforcing maximum penalties for possession and use, including for research purposes (Ninnemann et al., 2012).

Despite the consensus that those initial studies have several shortcomings regarding scientific or statistical rigor (Vollenweider and Kometer, 2010), they were the first to suggest the clinical use of these substances, which has been supported by recent data from both animal and human studies (Danforth et al., 2016; Nichols, 2004; Sartori and Singewald, 2019b). Moreover, some psychedelics are currently used as treatment options for psychiatric disorders. For instance, ketamine is prescriptible to treat TRD in USA and Israel, with many other countries implementing this treatment (Mathai et al., 2020), while Australia is the first nation to legalize the psilocybin for mental health issues such as mood disorders (Graham, 2023). Entactogen drugs such as the 3,4-Methylenedioxymethamphetamine (MDMA), are in the last stages of clinical research and might be employed for the treatment of post-traumatic stress disorder (PTSD) future studies on relevant disease models, such as depression, anxiety, and Alzheimer’s disease, must carefully consider such experimental parameters, including the age of the animals, treatment regimens, and time points of analysis.

with assisted psychotherapy (Emerson et al., 2014; Feduccia and Mithoefer, 2018; Sessa, 2017)

Nevertheless, as psychedelics edge closer to mainstream therapeutic use, we believe it is of utmost importance for mental health professionals to appreciate the role of set and setting in shaping the psychedelic experience (Hartogsohn, 2017). Drug developers, too, should carefully evaluate contraindications and potential interactions, given the unique pharmacological profiles of these compounds and the relative lack of familiarity with them within the clinical psychiatric practice. It would be advisable that practitioners intending to work with psychedelics undergo supervised clinical training and achieve professional certification. Such practical educational approach based on experience is akin to the practices upheld by Amerindian traditions, and are shown to be beneficial for treatment outcomes (Desmarchelier et al., 1996; Labate & Cavnar, 2014; Naranjo, 1979; Svobodny, 2014).

In summary, the rapidly evolving field of psychedelics in neuroscience is providing exciting opportunities for therapeutic intervention. However, it is crucial to approach this potential with due diligence, addressing the intricate balance of variables that contribute to the outcomes observed in pre-clinical models. The effects of psychedelics on neuroplasticity underline their potential utility in various neuropsychiatric conditions, but also stress the need for thorough understanding and careful handling. Such considerations will ensure the safe and efficacious deployment of these powerful tools for neuroplasticity in the therapeutic setting.

## Acknowledgements

RVLC thanks the colleagues from the Brain Institute and mentors from the PPG in Psychobiology (UFRN) for support and guidance, as well as CAPES (*Coordenação de Aperfeiçoamento de Pessoal de Nível Superior*, Brazil) for the scholarship funding. TCM is grateful for the funding from the Royal Physiographic Society of Lund and the Segerfalk Foundation.

