## Supplementary Tables 1-6 for "Effects of psychedelics on neurogenesis and brain plasticity: A systematic review"

*Supp Table 1 General methods used by reviewed CB1 agonists studies*

| Compound | Dose/Via | Frequency of treatment | Disease model | Animal model (# of subjects/group) | Type of neurogenesis | Reference |
|---|---|---|---|---|---|---|
| ***In vivo* studies** | | | | | | |
| **AEA** | AEA (2 µM); WIN 55, 212-2 (2 µM); | Acute (5-10 minutes) | No | Mice C56BL7, E13.5-E18.5, PN145-PN180 $CB1R^{f/f;\ Dlx5/6\text{-}Cre}$ mice (n= 5); Rat (n=ND) | Embryonic | (Berghuis et al., 2007) |
| **HU210** | HU210 (0.010, 0.050 mg/kg) IP | Chronic (10, 20 days) | AD(APP23/PS45) | Mice P28 and P63 (Total = 89) | Adult | (Chen et al., 2010) |
| **HU210** | HU210 (0.1 mg/kg) IP | Acute (twice daily); Chronic (10 days) | No | Rats male Long-Evans, Wistar, Fischer (n= 3-7) | Adult | (Jiang et al., 2005) |
| **HU210** | HU210 (0.025, 0.050, 0.100 mg/kg) IP | Chronic (11 days twice a day) Dose escalate in 3 age windows (0.025mg/kg in P35-P37; 0.050mg/kg in P38-P41; 0.100mg/kg in P42-P45) | No | Rats Sprague Dawley P28 (n= 5-6) | Adult | (Lee et al., 2014) |
| **M-AEA** | M-AEA (5mg/kg) IP | Chronic (4 days) | No | Rats Sprague-Dawley (n=4-8) | Adult | (Rueda et al., 2002) |
| **THC** | THC (3 mg/kg) IP | Acute | No | Mice CD-1 Female P35, P60, P90 (n=8) | Adult | (Leishman et al., 2018) |
| **THC** | THC (1, 10, 30 mg/kg) SC | Acute | No | Rats Male P4-P7, P20-P25, P90-P120 (n=4-9) | Developmental, Adult | (Downer et al., 2007) |
| **THC** | THC (2.5, 5, 10 mg/kg) Daily oral | Chronic (11 days twice a day) Dose escalate in 3 age windows (2.5mg/kg in P35-P37; 5mg/kg in P38-P41; 10mg/kg in P42-P45) | No | Rats Sprague-Dawley P28 (Total = 164) | Adult | (Rubino et al., 2008) |
| **THC** | THC (2.5, 5, 10 mg/kg) Daily oral | Chronic (11 days twice a day) Dose escalate in 3 age windows (2.5mg/kg in P35-P37.; 5mg/kg in P38-P41; 10mg/kg in P42-P45) | No | Rats Females Sprague-Dawley P28 (n=4-6) | Adult | (Realini et al., 2011) |
| **THC** | THC (6 mg/kg) IP | Chronic (15 days) | No | Rats Male Sprague-Dawley P28-42 (n= 10-11) | Adult | (Steel et al., 2014) |
| **THC** | THC (0.75, 1.5, 3 mg/kg) IP | Acute (for 7 days) Chronic (21 days) | No | Rats Male Sprague-Dawley P28 (n=6) | Adult | (Suliman et al., 2018) |

| | | | | | | |
|---|---|---|---|---|---|---|
| **THC** | THC (2.5, 5, 10 mg/kg) Daily oral | Chronic (11 days twice a day) Dose escalate in 3 age windows (2.5mg/kg in P35-P37; 5mg/kg in P38-P41; 10mg/kg in P42-P45) | No | Rats Female Sprague-Dawley P28 (n=4-8) | Adult | (Cuccurazzu et al., 2018) |
| **THC; Win 55212-2** | THC (1,3, 10, 30 mg/kg) IP;<br>20 to 80 mg/kg daily oral. | Acute; Repeated; Chronic (21 days) | No | Mice C57BL6 Male (n=4-9) | Adult | (Kochman et al., 2006) |
| **THC** | THC (1, 5 mg/kg) s.c. | Chronic (12 and 24 days) | No | Male Mice P5-P35 (n = 3-8) | Developmental, Adult | (Beiersdorf et al., 2020) |
| **THC** | THC (0.3, 1, 3 mg/kg) i.p. | Chronic (escalating 0.3 mg/kg P35-37 -1 mg/kg P38-41, 3 mg/kg P42-45) | No | Male Rats (n =7-12 behavioral experiments; n = 5 IHC experiments) | Adult | (Poulia et al., 2021) |
| ***In vitro* studies** | | | | | | |
| **AEA** | AEA (5 µM) | Acute (24h exposure) | No | Rats Cultured Cortical Neurons (n = 3 cultures); HNSC.100 (n = 3 cultures); PC12 cells (n = 3 cultures) | N/A | (Rueda et al., 2002) |
| **AEA; THC** | THC (1µM) ; AEA (1-10 µM) * | Chronic (21 days) * | No | hCBiPSC (n= 3-5 cultures) for Electrophysiological experiments (n= 2-35 cells) | N/A | (Stanslowsky et al., 2017) |
| **AEA; Win 55,212-2** | AEA (100 µM); Win55,212-2 (20 µM) | Acute (15-30 minutes) | No | Cultured GABAergic neurons from Mice E18.5 expressing CB1R (N = 13) ; Xenopus laevis (N = 8-210) | Embryonic | (Berghuis et al., 2007) |
| **HU210** | HU210 (1, 10, 100nM, 1, 10 µM) | Acute (48hrs exposition) Chronic (8 days) | No | Cultured Hippocampal NS/PCs cells derived from E17 (n= 4-6 cultures) | Embryonic | (Jiang et al., 2005) |
| **HU210** | HU210 (0.01, 0.1, 1. 10, 100 nM) | Acute (10-15 minutes exposure) | No | Mouse Neuroblastoma N1E-115 cells (n = 3 cultures) | N/A | (Zhou and Song, 2001) |
| **HU210** | HU210 (0.03, 0.3,3 µM) | Acute (6 days) | Hyperglycemic neuropathy | PC12 cells (n = 4-6 cultures; 136-218 cells) | N/A | (Zhang et al., 2009) |

| | | | | | | |
|---|---|---|---|---|---|---|
| **O-2545** | O-2545 (0.035,0.07, 0.35mg/ml) | Acute (10µl solution during gastrulation above the ventral side of embryos) | No | Chicken embryos S3$^{+}$-E8$^{-}$, Charles River lab (n=3-54) | Embryonic | (Psychoyos et al., 2008) |
| **THC** | THC (5nM - 50µM) | Acute (2h) | No | Rat Cultured Cortical Neurons (n = 5-6 cultures) | N/A | (Downer et al., 2007) |
| **THC** | THC (3 µM); 2-AG (1 µM) | Acute (15-30 minutes and 24 hours) | No | hiPSC (n= 3 cultures per hiPSC line) | N/A | (Shum et al., 2020) |
| **THC** | THC (100 nM, 1, 7.5, 10 µM) | Acute (24 hours) | No | mice E14.5 cortex culture (n =15-33) | N/A | (Beiersdorf et al., 2020) |
| **Human studies** | | | | | | |
| **THC; CBD** | THC/CBD ratio of hair samples and self-report of consumption | N/A | No | Healthy cannabis users and IQ, age matched controls (N=11-13) | N/A | (Demirakca et al., 2011) |

AD (Alzheimer's Disease); THC ((−)-trans-Δ9-tetrahydrocannabinol); N/A (Not Applicable); SC (Sub-Cutaneous); HNSC.100 (human neural stem cell line 100); PC12 (cell line derived from a pheochromocytoma of the rat adrenal medulla);* = complex dosage and time exposition due to elevated cytotoxicity on concentrations above 10 µM for both drugs; hCBiPSC (human cord blood-derived induced pluripotent stem cell); IQ (Intelligence quotient); NS/PCs (Neural Stem/Progenitor Cells); S# ( # of somites in the embryo).

*Supp Table 2. General methods used by reviewed NMDA antagonists' studies*

| Psychedelic Used | Dose/via | Frequency of treatment | Disease model | Animal model (# of subjects/group) | Type of Neurogenesis | Reference |
|---|---|---|---|---|---|---|
| | | | ***In vivo* Studies** | | | |
| **Ketamine** | Ketamine (30mg/kg) IP | Acute (3 days at P7, P9, P11) | No | Mice Male C57BL6 P11 and P70 (n = 4-7) | Developmental | (Schiavone et al., 2020) |
| **Ketamine** | Ketamine (30mg/kg) IP | Chronic (14 days) | Alcohol co-administration | Rats Male Sprague Dawley P35-42 (n=3-8) | Adult | (Zuo et al., 2018) |
| **Ketamine** | Ketamine (1, 10 mg/kg); Ketamine/lithium (1mg/10mg/kg) IP | Acute | No | Rats Male Sprague-Dawley (n= 4-8 animals; 8-17 cells) | Adult | (Liu et al., 2013) |
| **Ketamine** | Ketamine (30 mg/kg) IP | Chronic (5 days) | No | Rats Male Sprague-Dawley P56 (n = ND) | Adult | (Keilhoff et al., 2004) |
| **Ketamine** | Ketamine (4, 8, 20 mg/kg) IP | Acute | No | Rats Male Sprague-Dawley P70-P95 (n = 7-14) | Adult | (Goulart et al., 2010) |
| **Ketamine** | Ketamine (5, 10, 80 mg/kg) IP | Acute | No | Rats Male Sprague-Dawley (n =4-8) | Adult | (Li et al., 2010) |
| **Ketamine** | Ketamine (0.25, 0.5 mg/kg); IP | Chronic (11 days) | No | Rats Male Wistar (n=8) | Adult | (Akinfiresoye and Tizabi, 2013) |
| **Ketamine** | Ketamine (10 mg/kg) IP | Acute | No | Rats Male Sprague Dawley (n = 6-8) | Adult | (Lepack et al., 2015) |
| **Ketamine** | Ketamine (40 mg/kg) IP | Acute (4 times with 1h inter-dose interval) | No | Rats P7 (n = 5-6) | Developmental | (Huang et al., 2016) |
| **Ketamine** | Ketamine (10 mg/kg) IP | Acute | No | Rats Females Sprague-Dawley P56 (n = 3 animals; 11-42 neurons | Adult | (Ly et al., 2018) |
| **PCP** | PCP (10 mg/kg/day) SC | Chronic (12 days from E6-E18) | No | Mice CD-1 P1/P7/P56 (n=5-20) | Developmental | (Toriumi et al., 2012) |
| **PCP** | PCP (7.5 mg/kg) IP | Acute; Chronic (5 and 14 days); Chronic with challenging Dose on day 21 | No | Rats Male Sprague-Dawley (n= 6) | Adult | (Liu et al., 2006) |
| **PCP** | PCP (5 mg/kg) SC | Chronic (14 days E8-E21) | No | Rats Sprague-Dawley P21 (n=3-8) | Embryonic | (Tanimura et al., 2009) |

| | | | | | | |
|---|---|---|---|---|---|---|
| **PCP** | PCP (1-10 mg/kg) IP | Chronic (14 days) | No | Mice Male CD-1 P56 (n = 6-7) | Adult | (Maeda et al., 2007) |
| | | | ***In vitro* Studies** | | | |
| **PCP** | PCP (1 µM) | Acute (24 hours) | No | Primary PFC mice cultured neurons (n = 3) | Developmental | (Zhang et al., 2016) |
| | | | **Human Studies** | | | |
| **Ketamine** | Ketamine dependence (PANSS questionnaire) | N/A | Hospitalized addicted subjects | Ketamine addicted (n= 155) and matching healthy control (n=80) | N/A | (Fan et al., 2015) |
| **Ketamine** | 0.71 mg/kg via ND | Acute | Cocaine non-treatment seeking addicted voluntaries | Cocaine addicted (n=20) | N/A | (Dakwar et al., 2018) |
| **Ketamine** | 0.5mg/kg IV | Acute (7 days after discontinuation of standard medication) | Bipolar patients within a depression episode and resistant to classical antidepressants | Treatment resistant depressive patients (n = 25) | N/A | (Rybakowski et al., 2013) |
| **Ketamine** | 0.5mg/kg IV | Acute | Treatment resistant MDD | Treatment resistant MDD (n = 30) | N/A | (Duncan et al., 2013) |
| **Ketamine** | 0.5mg/kg IV | Acute | Treatment resistant MDD | Treatment resistant MDD (n=22) | N/A | (Haile et al., 2014) |

ND (Not Described); IP (Intraperitoneal); IV (Intravenous); SC (Subcutaneous); N/A (Not Applicable); P# (Post-Natal Day #); E# (Embryonic day #); MDD (Major Depressive Disorder); PFC (Pre-Frontal Cortex); PANSS (Positive and Negative Syndrome Scale).

*Supp Table 3. General methods used by reviewed Harmala alkaloids studies*

| Psychedelic used | Dose/Via | Frequency of treatment | Disease model | Animal model (# of subjects/group) | Type of neurogenesis | Reference |
|---|---|---|---|---|---|---|
| | | | ***In vivo* studies** | | | |
| **Harmine** | Harmine (5, 10, 15 mg/kg) IP | Acute | No | Rats Male Wistar P60 (n=15) | Adult | (Fortunato et al., 2009) |
| **Harmine** | Harmine (5, 10, 15 mg/kg) IP | Chronic (14 days) | No | Rats Male Wistar P60 (n=15) | Adult | (Fortunato et al., 2010a) |
| **Harmine** | Harmine (15mg/kg) IP | Chronic (7 days) | Chronic Mild Stress | Rats Male Wistar P60 (n=15) | Adult | (Fortunato et al., 2010b) |
| **Harmine** | Harmine (10, 20 mg/kg) IP | Chronic (10 days) | Unpredictable Chronic Stress | Mice C57BL6 Male P56-P70 (n=10) | Adult | (Liu et al., 2017) |
| | | | ***In vitro* studies** | | | |
| **Harmine** | Harmine (7.5 μM) | Not reported | No | hNPCs (n = 6) | N/A | (Dakic et al., 2016b) |
| **Harmine; Tetrahydroharmine; Harmaline; Harmol** | Harmine (1 μM); T-Harmine (1 μM); Harmaline (1 μM); Harmol (1 μM) | Chronic (7 days); Acute (3 days) | No | Neurospheres from adult mice SGZ and SVZ (n= 3 independent cell cultures) | Adult | (Morales-García et al., 2017) |

SVZ (Subventricular Zone); SGZ (Subgranular Zone); IP (Intraperitoneal); hNPCS (Human neural progenitor cells).

*Supp Table 4 General methods used by reviewed Psychoactive tryptamines studies*

| Psychedelic used | Dose/Via | Frequency of treatment | Disease model | Animal model (# of subjects/group) | Type of neurogenesis | Reference |
|---|---|---|---|---|---|---|
| | | | ***In vivo* studies** | | | |
| **DMT; LSD; DOI; Ketamine.** | Drosophila experiments, DOI, LSD (ND);<br>DMT 1, 10 mg/kg<br>Ketamine 10 mg/kg | Acute | No | Drosophila (n= ND); Rats Females Sprague-Dawley P56 (n = 3 animals; 11-42 neurons) | Adult | (Ly et al., 2018) |
| **5-MeO-DMT** | 5-MeO-DMT (100 µg) ICV | Acute | No | Mice C56BL7 P55-P70 (n =3-6) | Adult | (Lima da Cruz et al., 2018) |
| **5-MeO-DMT** | 5-MeO-DMT (20mg/kg) IP. | Acute | Tinnitus (acute salicylate model) | Mice C56BL7 P120-P150 (N=8) | Adult | (Winne et al., 2020) |
| **Ibogaine** | Ibogaine (20, 40 mg/kg) IP. | Acute | No | Rats Male Wistar P (n = 6) | Adult | (Marton et al., 2019) |
| **Methylpsilocin** | Methylpsilocin (0.7 mg/kg) IP | Acute and sub-acute (7 days) | CUS | Rats Male Wistar (N=6) | Adult | (Wankhar et al., 2020) |
| **BW723C86; MDMA, DOI,** | BW723C86 (1µm); MDMA (20 mg/kg) IP.; DOI (1,3,5mg/kg) IP. | Acute | 5-HT2B KO; WT | Mice Male mixed B6::129S2 P21-28 (n= 3-5) | Adult | (Belmer et al., 2018) |
| **Psilocybin; NBOMe** | Psilocybin (0.1,0.5,1, 1.5 mg/kg) IP.; NBOMe (0.1,0.3,1 mg/kg) IP | Acute | No | Mice male C57BL6 (n=6) | Adult | (Catlow et al., 2013) |
| **Ayahuasca** | Ayahuasca (250, 500, 800 mg/kg) oral | Acute | No | Rats Male Wistar (n= 5-8) | Adult | (Castro-Neto et al., 2013) |
| **DMT and many isoforms** | mice = DMT (5ml/kg*) IP.; Zebrafish = DMT (28µM-200µM) | Acute | No | Mice both genders C57BL6 P (N= 3-8); Zebrafish larvae (n = 8 larvae per well, 18 wells per conditions | Adult/Developmental | (Dunlap et al., 2020) |
| **DMT** | DMT (2mg/kg) i.p. | Acute (4 days); Chronic (every other day for 21 days) | No | Adult Male Mice P90 (n = 5) | Adult | (Morales-Garcia et al., 2020) |

| | | | | | | |
|---|---|---|---|---|---|---|
| | | | *In vitro* **studies** | | | |
| **Ayahuasca** | all compounds tested at 1, 1.5, 2.5, 10.5 µg/mL | Acute (48h-72h incubation) | 6-OHDA induced Neurodegeneration (Parkinson's Disease model) | SH-SY5Y cells (n= 3 cultures) | N/A | (Katchborian-Neto et al., 2020) |
| **DMT and many isoforms** | DMT (10µM -10 pM); IsoDMT (10µM -10 pM) | Acute (1h exposure) | No | Rats Cultured Cortical Neurons 6 days-in-vitro (n=46-79 neurons) | N/A | (Dunlap et al., 2020) |
| **DOI; DMT; LSD** | DOI (10 µM); DMT (90 µM); LSD (10 µM). | Acute (24h exposure) | No | Rats Cultured Cortical Neurons 19 days-in-vitro (n=46-79 neurons) | N/A | (Ly et al., 2018) |
| **DMT** | DMT (1µM) | Acute (24 hours); Chronic (7 days) | No | Culture mice SGZ cells (n =3 independent cell cultures) | N/A | (Morales-Garcia et al., 2020) |

IP (intraperitoneal); ICV (intracerebroventricular); N/D (Not Declared); N/A (Not Applicable); CUS (Chronic Unpredictable Stress model); MP (methylpsilocin) SH-SY5Y (human cell line derived from neuroblastoma).

*Supp Table 5 General methods used by Entactogens reviewed studies*

| Psychedelic used | Dose/Via | Frequency of treatment | Disease model | Animal model (# of subjects/group) | Type of neurogenesis | Reference |
|---|---|---|---|---|---|---|
| | | | **In vivo** | | | |
| **MDMA** | MDMA (1.25, 20mg/kg) Oral | Chronic (36 days) | No | Mice from embryo to adolescence (E6-P21) C57BL6 (n=16) | Embryonic/Adult | (Cho et al., 2008) |
| **MDMA** | MDMA (10mg/kg) IP. | Chronic (4x daily with a 2 h interdose interval by 10 days) | No | Rats Male Sprague Dawley P11-20 (n= 16-27) | Developmental | (Schaefer et al., 2013) |
| **MDMA** | MDMA (5.mg/kg) IP. | Acute (8 times with a 6h interdose interval) | No | Rats Male Wistar (n= 6-7) | Adult | (Hernández-Rabaza et al., 2006) |
| **MDMA** | MDMA (1.25, 2.5, 5 mg/kg) IP. | Chronic (10 days) | No | Rats Male Sprague-Dawley P28-39 (n=7-11) | Adult | (Catlow et al., 2010) |
| **MDMA** | MDMA (15mg/kg) Oral | Chronic E14-E20 (2x at day with an 8h interdose interval) | No | Rats Sprague-Dawley E14-P21 (n = 6) | Embryonic | (Thompson et al., 2012) |
| **MDMA** | MDMA (10mg/kg) Oral | Acute E13-E15 (2x at day with an 8h interdose interval) | No | Rats Long Evans E13-E15(n = 6) | Embryonic | (Canales and Ferrer-Donato, 2014) |
| **MDMA** | MDMA (5.mg/kg) IP. | Acute (3x at day with 2-3h interdose interval for 1 day) or<br>Chronic (3x at day with 2-3h interdose interval for 4 days) | No | Rats Male Sprague-Dawley P27-P29/P48-P50 (n= 18) | Adult | (García-Cabrerizo and García-Fuster, 2016) |

IP (intraperitoneal).

*Supp Table 6 General methods used by reviewed studies using other psychedelics.*

| **Psychedelic used** | **Dose/Via** | **Frequency of treatment** | **Disease model** | **Animal model (# of subjects/group)** | **Type of neurogenesis** | **Reference** |
|---|---|---|---|---|---|---|
| | | | ***In vivo* studies** | | | |
| **DOI** | DOI (1μm) | Acute | 5-s$HT_{2A}$R-KO | Mice "WT" and 5-$HT_{2A}$R-kO P14-P21 (n = ND, cells = 5-13) | Adult | (Berthoux et al., 2019) |
| **DOI; LSD** | DOI (8mg/kg); LSD (0.5mg/kg) IP. | Acute and chronic (7 days), same doses | No | Rats Male Wistar (n=4 - 7) | Adult | (Jha et al., 2008) |
| | | | ***In vitro* studies** | | | |
| **DOI** | DOI (10, 100, 500 nM, 1 μM) | Acute (5-60 minutes) | No | Cultured rat cortical neuron (n = ND); Cos-7 cells (n=ND) | N/A | (Jones et al., 2009) |
| **DOI** | DOI (1, 2, 5 μM) | Acute (30min or 120 min exposure); Chronic (6 days exposure) | No | Human neuroblastoma SK-N-SH cells (n=3 independent cell cultures) | N/A | (Marinova et al., 2017) |

WT (Wild Type); IP (Intraperitoneal); Cos-7 (); ND (Not described); N/A (Not Applicable).
